# Hepatobiliary Progenitor-like Reprogramming in Liver Metastases

**DOI:** 10.64898/2026.06.30.735617

**Authors:** Ajit Kumar Sharma, Nobuyuki Takahashi, Yingying Cao, Abhinav Joshi, Sophie Zhuang, Amira Kazi, Michael Nirula, Sarthak Sahoo, Yang Zhang, Rajesh Kumar, Seemadri Subhadarshini, Kanak Parmar, Melissa Mikolaj, Roshan L. Shreshta, Tiyam Assadpour, George Chrisafis, Parth Desai, Asrar Alahmadi, Hui-Zi Chen, Samantha Nicholas, Yue Huang, Maria Sebastian Thomas, Ross Lake, Neel Sanghvi, Nishanth Nair, Nivedita Mukherjee, Bhavya Dhaka, Christopher A. Febres Aldana, Sabarinathan Radhakrishnan, Nir Friedman, G. Tom Brown, David E. Kleiner, Simone Difilippantonio, Dwight H. Owen, Thorkell Andersson, Eytan Ruppin, Kedar Narayan, Mohit Kumar Jolly, Stephen Hewitt, Anish Thomas

## Abstract

Metastatic progression requires cancer cells to adapt to the unique constraints of distant organ microenvironments, yet the mechanisms that drive organ-specific adaptations remain poorly understood. Here, we show that the liver actively rewrites metastatic cancer cell identity, driving tumor cells toward a hepatobiliary progenitor-like state. Through integrated transcriptomic, proteomic, metabolomic, and epigenomic analyses of patient-derived rapid-autopsy samples and experimental models, we identify this state as selectively enriched in liver metastases. It is characterized by co-activation of hepatic and biliary/progenitor regulators HNF4A and SOX9 and is observed across multiple epithelial cancers, indicating a conserved response to the hepatic niche. Mechanistically, hepatocyte-derived TGF-β and hypoxia converge to activate a HIF-1α-ACLY axis, increasing nuclear acetyl-coenzyme A availability and histone acetylation at hepatic lineage regulatory elements to drive hepatobiliary reprogramming. This coordinated niche-response program can be captured transcriptionally and is associated with inferior overall survival. Disruption of this pathway suppresses hepatic reprogramming and impairs liver metastatic fitness. These findings identify the liver as an active determinant of metastatic cell fate, linking microenvironmental signaling to metabolic and chromatin remodeling programs that enable lineage plasticity. More broadly, they reveal organ-specific reprogramming as a fundamental principle of metastasis and a therapeutic vulnerability in liver metastases.

## Introduction

Metastasis causes the vast majority of cancer deaths across solid tumors^1,2^, yet the mechanisms that enable disseminated cancer cells to colonize and adapt to distinct organ microenvironments remain incompletely understood^3^. The liver is among the most common and clinically consequential sites of metastasis in lung, colorectal, pancreatic, breast, and other cancers^4^. Hepatic involvement is associated with rapid clinical decline and resistance to therapy^5^, but the molecular programs that enable tumor cells to survive and expand within the liver remain poorly defined.

Metastatic adaptation often involves the co-option of developmental and regenerative programs normally deployed during tissue injury and repair^6^. These programs can reactivate lineage-defining transcription factors, promote cell-state transitions, and remodel chromatin accessibility^7–11^, frequently without metastasis-specific genetic driver alterations^12,13^. In parallel, distinct metastatic organs impose unique metabolic constraints, revealing dependencies shaped by nutrient availability, tissue architecture, oxygen tension, and stromal signaling^14^. How these extrinsic niche cues converge on tumor-intrinsic regulatory circuits to rewire metastatic cell identity remains a central unanswered question.

Small cell lung cancer (SCLC) provides a powerful model to investigate organ-specific metastatic adaptation^15,16^. This highly aggressive neuroendocrine carcinoma is defined by early dissemination, frequent liver metastasis, lineage plasticity, rapid proliferation, and profound therapeutic resistance^17–21^. Patients with SCLC liver metastases have particularly poor outcomes, yet the biology of liver colonization has been difficult to study because metastatic tissues are rarely sampled and few experimental systems capture organ-specific tropism and adaptation^22^.

Here, we integrate transcriptomic, proteomic, metabolomic, and epigenomic profiling of liver and non-liver metastases from patients with SCLC, including biopsy and rapid-autopsy specimens, with functional modeling *in vitro* and *in vivo*. We identify a hepatobiliary progenitor-like state selectively enriched in SCLC liver metastases that mirrors liver regenerative programs. Mechanistically, hepatocyte-derived TGF-β and hypoxia converge on a HIF-1α–ACLY metabolic axis to increase nuclear acetyl–coenzyme A availability, remodel the chromatin landscape, and reshape metastatic cell identity. A composite NichePolarity score integrating hepatobiliary progenitor features with TGF-β and hypoxia programs capture this liver-adapted transcriptional state, which is associated with inferior overall survival. Together, these findings establish the liver as an active regulator of metastatic cell fate and reveal organ-specific lineage reprogramming as a source of therapeutic vulnerability in liver metastases.

## Results

### SCLC liver metastases are clinically aggressive and acquire liver-associated features

The liver is a frequent site of metastasis across solid tumors and is associated with poor clinical outcomes^23^. In SCLC, liver involvement is common but poorly understood, in part because metastatic tissues are rarely available for molecular analysis. We assembled a cohort of 105 SCLC tumors, including 38 liver metastases, collected via clinical biopsy or rapid autopsy (**Fig. 1A, fig. S1A, table S1**). Patients with liver metastases had significantly shorter progression-free survival following platinum-based chemotherapy (median, 60 days vs. 90 days; HR = 2.16; P = 0.030) and reduced overall survival (median, 12.0 vs. 16.2 months; HR = 2.47; P = 0.0007; **Fig. 1B**). Liver metastasis remained independently associated with poor survival in multivariable analysis (**table S2**).

**Fig. 1:**
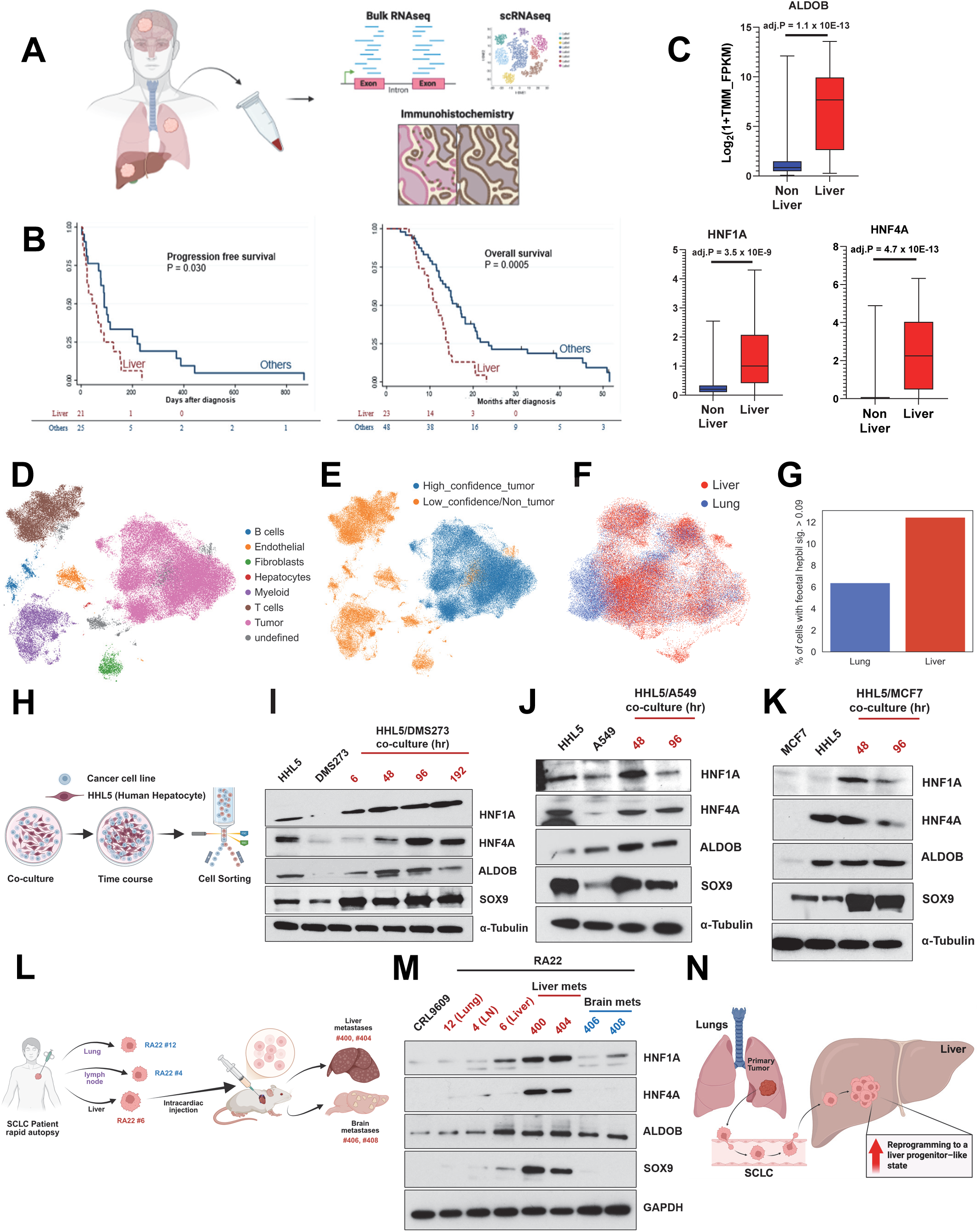
SCLC liver metastases are clinically aggressive and undergo liver progenitor-like reprogramming. **(A)** Schematic overview of the multi-omic strategy integrating bulk and single-cell RNA-seq and IHC to profile liver and non-liver SCLC metastases. **(B)** Kaplan-Meier analysis showing progression-free survival and overall survival in patients with liver metastases (red dotted line, n = 38) compared to those with non-liver metastases (blue line, n = 67); P = 0.030 and P = 0.0005 by log-rank test. **(C)** Expression of representative hepatocyte-associated genes in liver (n = 38) versus non-liver (n = 67) metastases based on bulk RNA-seq (log2 TPM values; adjusted P-values shown). **(D)** UMAP embedding of integrated scRNA-seq data showing broad cell-type categorization across pooled SCLC patient tumor samples (n = 39). **(E)** RB1 loss-based classification of malignant versus non-malignant cells. Cells with high RB1-loss signature scores were annotated as high-confidence tumor (blue) and others as low-confidence/non-tumor (orange). **(F)** UMAP visualization highlighting cells from liver metastases (red) and lung (blue) tumors. **(G)** Quantification of the proportion of cells exceeding the 90th percentile fetal hepatobiliary progenitor signature threshold in lung vs. liver metastases. The full distribution of signature scores is shown in Fig. S1F. **(H)** Experimental design of the hepatocyte co-culture system, in which cancer cell lines were co-cultured with human hepatocytes (HHL5) followed by time-course sampling and fluorescence-based sorting to isolate cancer cells for downstream analyses. **(I)** Immunoblot analysis of DMS273 SCLC cells co-cultured with hepatocytes for the indicated time points (6–96 h) showing induction of hepatocyte-lineage transcription factors HNF4A and HNF1A, the progenitor-associated SOX9, and the liver-enriched metabolic enzyme ALDOB. (**J–K**) Immunoblot validation of liver progenitor-like reprogramming in non-SCLC cancer cell lines (A549 lung adenocarcinoma and MCF7 breast cancer) following co-culture with hepatocytes. **(L)** Schematic of metastatic SCLC cell-line model generation from rapid autopsy specimens. Tumor-derived cell lines established from lung (RA22 #12), lymph node (RA22 #4), and liver (RA22 #6) metastases were expanded and injected intracardially into immunodeficient mice to generate organ-specific metastatic derivatives, including liver-derived lines (#400, #404) and brain-derived lines (#406, #408). **(M)** Immunoblot validation of liver progenitor-like reprogramming in P0_liver and P1_liver lines compared to P1_brain, lymph node, lung, or normal bronchial epithelial controls (CRL9609). **(N)** Conceptual model illustrating liver progenitor-like reprogramming in liver metastasis. **Abbreviations:** SCLC, small cell lung cancer; scRNA-seq, single-cell RNA sequencing; UMAP, Uniform Manifold Approximation and Projection; PCA, principal component analysis; IHC, immunohistochemistry; HR, hazard ratio.

To define molecular features of liver-metastatic SCLC, we performed bulk RNA sequencing on liver (n = 38) and non-liver (n = 67) tumors. Liver metastases showed marked up-regulation of hepatocyte-associated genes, including *HNF1A*, *ALDOB*, and *HNF4A* (**Fig. 1C**). Pathway analysis demonstrated enrichment of liver metabolic programs, including xenobiotic metabolism, bile acid synthesis, and fatty acid oxidation, together with reduced immune and inflammatory signaling (**fig. S1B**). These findings were reproduced in a patient-matched rapid-autopsy cohort comprising 32 tumors from five patients, including liver and non-liver metastases from the same individuals (**fig. S1B, table S3**)^24^. Somatic mutation frequencies did not differ significantly between liver and non-liver sites (**fig. S1D**), suggesting that the liver-associated transcriptional program was not explained by recurrent site-specific genetic alterations.

We next performed proteomic profiling of laser-capture microdissected tumor regions from 10 patients across metastatic sites. Consistent with the transcriptomic data, liver metastases were enriched for proteins involved in xenobiotic metabolism, bile acid synthesis, and fatty acid oxidation (**table S4, fig. S1E**). Thus, SCLC liver metastases are associated with adverse clinical outcomes and acquire transcriptional and proteomic features of the hepatic microenvironment.

### Activation of liver-progenitor-like features in SCLC liver metastases

We next asked whether the liver-associated program reflected tumor-intrinsic reprogramming rather than contamination by hepatic parenchyma. We analyzed single-cell RNA sequencing data from newly generated and publicly available SCLC tumor samples (n = 39; **table S5**) ^25,26^. After integration and batch correction, clustering resolved major immune, stromal, and epithelial/tumor compartments defined by canonical lineage markers (**Fig. 1D**).

To distinguish malignant from non-malignant epithelial cells, we leveraged RB1 loss, a defining genomic feature of SCLC^27,28^. Each cell was scored using a validated *RB1*-loss gene signature^29^, and cells exceeding a conservative threshold (score > 0.025) were classified as high-confidence tumor. This approach effectively delineated the tumor cells while excluding immune and stromal cells (**Fig. 1E**). As an internal control, *RB1*-loss-upregulated genes were selectively enriched in the tumor cluster (**fig. S1D**), while a reference hepatocyte signature scored highest in bona fide hepatocytes and remained low in immune and stromal cells (**fig. S1E**) supporting the specificity of both signatures.

Restricting analysis to these high-confidence tumor cells (**Fig. 1F**), we assessed activation of developmental hepatobiliary programs using a fetal hepatobiliary hybrid progenitor (HHyP) signature^30^. Tumor cells from liver metastases showed increased fetal HHyP signature scores relative to lung-derived tumor cells (**fig. S1F**). Using a 90th-percentile threshold to define high-scoring cells (**fig. S1F, red horizontal line**), the proportion of HHyP-high tumor cells was approximately doubled in liver metastases compared with lung lesions (12.5% versus 6.3%; **Fig. 1G**). This enrichment was observed across multiple patients, was not driven by a single outlier, and remained consistent across alternative thresholds (e.g., 85th or 95th percentile) and after exclusion of undefined clusters. These data indicate that a subset of SCLC cells in liver metastases acquire a tumor-intrinsic hepatobiliary progenitor-like state.

We validated this program at the protein level in matched liver and non-liver metastases from three rapid-autopsy cases. Immunohistochemistry for ALDOB, a liver-enriched metabolic enzyme regulated by HNF1A and HNF4A, showed strong cytoplasmic staining in liver metastases but minimal or absent staining in patient-matched non-liver sites, including lymph node, lung, adrenal, pancreas, and pericardium (**fig. S1G**). Adjacent hepatocytes stained strongly positive for ALDOB, serving as internal positive controls. Notably, ALDOB expression was concentrated at the tumor–liver interface and was largely absent from the tumor core (**fig. S1H**), consistent with a spatially restricted, microenvironment-dependent reprogramming event. Together, these findings show that SCLC liver metastases activate a hepatobiliary progenitor-like program in tumor cells, most prominently at sites of direct engagement with the hepatic niche.

### Experimental models recapitulate liver-progenitor like reprogramming

During liver regeneration, mature HNF4A-positive hepatocytes can transiently dedifferentiate into bipotent progenitor-like states marked by SOX9 and other progenitor-associated regulators^31–34^. These cells retain the ability to give rise to both hepatocytes and cholangiocytes, enabling restoration of tissue integrity following liver injury. Because SCLC liver metastases activated fetal HHyP programs in patient-derived SCLC liver metastases, we hypothesized that hepatic niche signals reprogram tumor cells toward a liver progenitor-like state.

To test this hypothesis, we developed an *in vitro* co-culture system in which GFP-labeled human SCLC cells were co-cultured with RFP-labeled human hepatocytes (**Fig. 1H, fig. S2A**). Co-culture induced a time-dependent increase in liver progenitor-associated markers in SCLC cells (**Fig. 1I**). HNF4A and SOX9 were rapidly induced, HNF1A showed modest upregulation, while ALDOB was induced at later time points. Similar induction occurred in an indirect transwell system, indicating that soluble paracrine factors were sufficient to initiate reprogramming (**fig. S2B**). The phenotype was reproduced in an independent SCLC cell line, H69 (**fig. S2C**). Upon removal from co-culture, marker expression returned to baseline within 72 hours (**fig. S2D**), indicating that this state depends on sustained exposure to hepatic microenvironmental cues, consistent with the spatially restricted expression patterns observed in patient tumors (Fig. S1F, G).

RNA sequencing of GFP-positive tumor cells isolated after 6 hours of co-culture revealed early upregulation of progenitor-associated genes, including SPP1, CD44, SOX9, and SOX4 (**fig. S2E**). Gene set enrichment analysis (GSEA) demonstrated significant enrichment of the fetal HHyP, and additionally pathways associated with hepatic repair, including Wnt and TGF-β signaling^30,31,34^, and hypoxia-responsive programs (**fig. S2F)**, suggesting coordinated activation of injury-regeneration pathways by the hepatic niche.

Similar reprogramming was observed across epithelial cancers. Co-culture of non–small-cell lung, breast, and colorectal cancer cell lines with hepatocytes induced HNF1A, HNF4A, ALDOB, and SOX9 expression in a time-dependent manner (**Fig. 1J, K, fig. S2G**). Thus, hepatobiliary progenitor-like reprogramming is not restricted to SCLC but represents a conserved response of metastatic cancer cells to hepatocyte-derived cues.

To assess reprogramming in vivo, we established an experimental metastasis model using rapid autopsy–derived SCLC cells (RA22; **Fig. 1L**). Intracardiac injection into NSG mice bypassed early metastatic steps such as vascular invasion, enabling focused interrogation of tumor-cell adaptation after colonization of distinct organ microenvironments^35^. This approach generated parental liver-derived cells (P0_liver) and matched metastatic derivatives from liver (P1_liver) and brain (P1_brain). Transcriptomic and proteomic profiling showed that liver-derived metastatic cells diverged from both parental and brain-derived cells (**fig. S2H, I**), with significant enrichment of fetal HHyP programs in liver-derived cells (**fig. S2J**). At the protein level, hepatic and progenitor markers were selectively enriched in liver-derived metastatic lines relative to parental cells, non-liver metastatic derivatives, lung- and lymph node-derived tumor cells, and normal bronchial epithelium (**Fig. 1M**). Together, these patient-derived and experimental data demonstrate that the hepatic niche induces a spatially restricted hepatobiliary progenitor-like state that is distinct from tumor states arising in non-hepatic microenvironments (**Fig. 1N**).

### Hepatic niche-derived TGF-β drives progenitor-like reprogramming in liver metastases

Our patient and model data suggested that metastatic SCLC cells acquire a hepatobiliary progenitor-like state in response to cues from the hepatic microenvironment. To test whether this state could be induced in tumor cells of non-hepatic origin, we introduced GFP-labeled DMS273 cells, originally derived from a malignant pleural effusion, into NSG mice by intracardiac injection. This generated liver metastases from which GFP-positive tumor cells were isolated for molecular profiling (**Fig. 2A, fig. S3A**). Liver-adapted DMS273 cells diverged transcriptionally from their parental counterparts (**fig. S3B**) and induced HNF1A, HNF4A, ALDOB, and SOX9 at the protein level (**Fig. 2A**), closely recapitulating the liver-associated state observed in patient-derived metastases and rapid-autopsy models. Similarly, intrasplenic injection of a patient-derived SCLC line isolated from lymph node metastasis produced liver lesions with a comparable hepatic shift (**fig. S3C**). Thus, hepatobiliary progenitor-like reprogramming is a reproducible consequence of hepatic colonization across models, routes of dissemination, and tumor cells of non-hepatic origin.

**Fig. 2:**
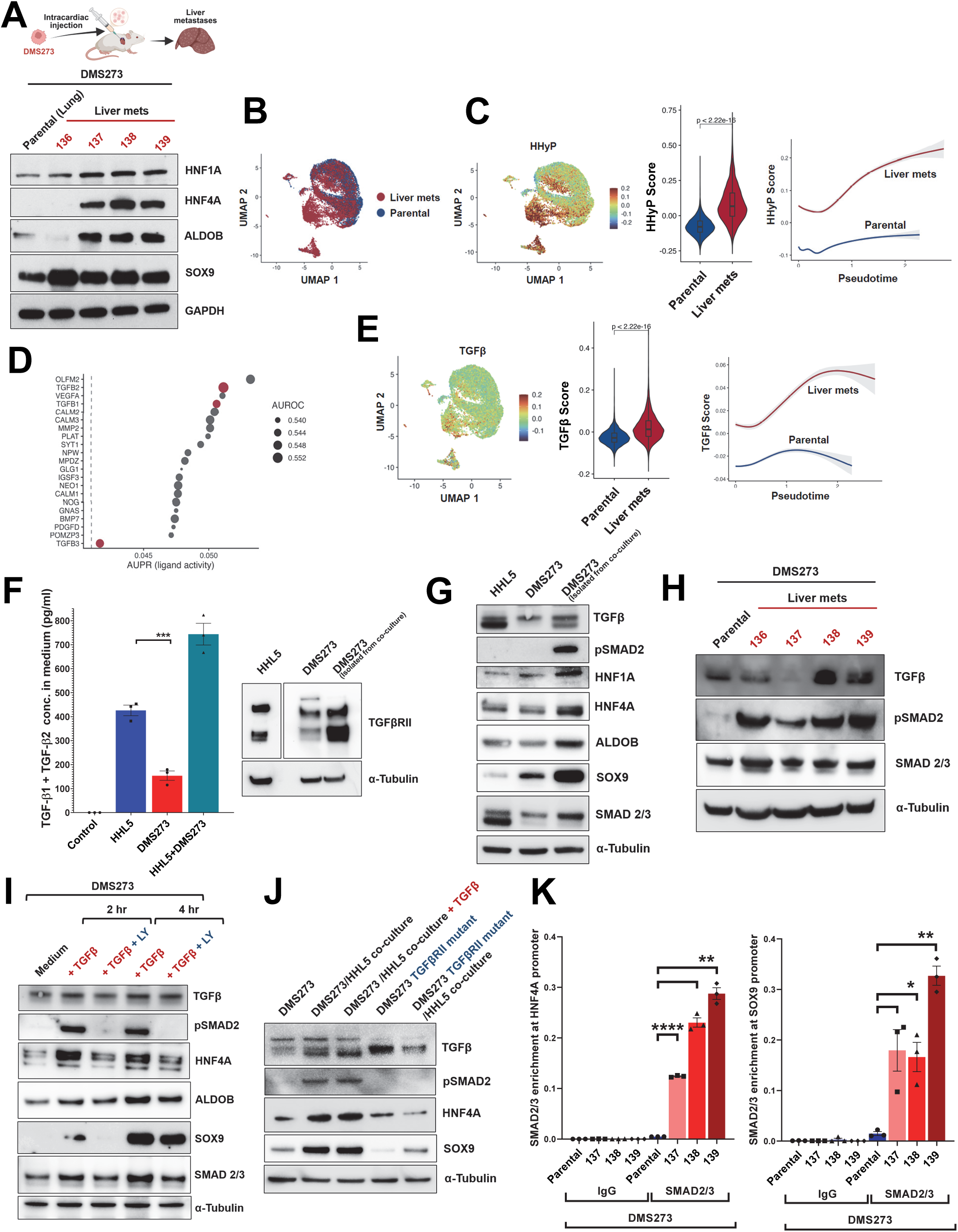
Hepatocyte-derived TGF-β instructs reprogramming in liver-metastatic SCLC. **(A)** Top, schematic of intracardiac injection of pleural effusion-derived DMS273 SCLC cells into NSG mice to generate liver metastases. Immunoblot analysis of matched parental and liver-metastatic derivatives for HNF1A, HNF4A, ALDOB, and SOX9. **(B)** UMAP projection of single-cell RNA-seq profiles from parental and liver-metastatic DMS273 tumors. **(C)** UMAP projection, module score distributions, and pseudotime analysis of fetal hepatobiliary progenitor (HHyP) program activity in parental and liver-metastatic cells. **(D)** Ligand-activity inference comparing liver-metastatic and parental tumor cells to identify candidate upstream regulators of the liver-adapted progenitor-like state, highlighting TGF-β family ligands. **(E)** UMAP projection, module score distributions, and pseudotime analysis of TGF-β pathway activity in parental and liver-metastatic cells. **(F)** Left, ELISA quantification of TGF-β1 and TGF-β2 in control medium and conditioned media derived from HHL5 hepatocytes, DMS273 cells, and HHL5–DMS273 co-culture (mean ± SEM, n = 3). Right, immunoblot analysis of TGF-β receptor II (TGF-βRII) expression in DMS273 cells following hepatocyte co-culture. **(G)** Immunoblot analysis of DMS273 cells cultured alone or with HHL5, probed for TGF-β, phospho-SMAD2, HNF1A, HNF4A, ALDOB, SOX9, and total SMAD2/3. **(H)** Immunoblot analysis of parental and *in vivo*-selected liver-metastatic DMS273 derivatives for TGF-β, phospho-SMAD2, and total SMAD2/3. **(I)** Immunoblot analysis of DMS273 cells treated with TGF-β in the presence or absence of the TGF-β inhibitor LY-3200882 for the indicated time points, probed for phospho-SMAD2, HNF4A, ALDOB, SOX9, and total SMAD2/3. **(J)** Immunoblot analysis of TGF-β receptor II mutant (TGFBR2-mutant) DMS273 cells cultured with or without HHL5 hepatocytes, probed for TGF-β, phospho-SMAD2, HNF4A, and SOX9. **(K)** ChIP-qPCR analysis of SMAD2/3 occupancy at regulatory regions of HNF4A and SOX9 in parental and liver-metastatic DMS273 cells (mean ± SEM, n = 3). Numbers 137, 138, 139 denote independent liver-metastatic derivatives obtained from separate animals. **Abbreviations:** SCLC, small-cell lung cancer; scRNA-seq, single-cell RNA sequencing; UMAP, Uniform Manifold Approximation and Projection; ChIP-qPCR, chromatin immunoprecipitation followed by quantitative PCR. Statistical significance: Statistical significance was assessed using two-sided Wilcoxon rank-sum tests for comparisons between groups or unpaired two-tailed *t*-test; *P* < 0.05 (*), < 0.01 (**), < 0.001 (***), < 0.0001 (****); ns = not significant.

To define this state at single-cell resolution, we performed single-cell RNA sequencing of parental and liver-metastatic DMS273 cells. Liver-derived cells formed a transcriptional state distinct from parental cells (**Fig. 2B**). Module scoring revealed increased fetal HHyP activity in liver-metastatic cells (**Fig. 2C**), and pseudotime reconstruction showed progressive enrichment of this program along liver-adapted trajectories. In contrast, parental cells showed low HHyP activity, consistent with a dynamic reprogramming process induced during liver adaptation.

To identify upstream signals driving this transition, we performed ligand-activity inference comparing liver-metastatic and parental tumor cells. TGF-β1 emerged as the top-ranked candidate ligand, with TGF-β2 and TGF-β3 also among the highest-scoring regulators (**Fig. 2D**). Among non TGF-β ligands, VEGFA was highly ranked, consistent with hypoxia and vascular remodeling within the liver metastatic niche^36^. Concordant with ligand inference, TGF-β pathway activity was increased in liver-metastatic cells relative to parental cells and rose progressively along liver-adapted pseudotime trajectories (**Fig. 2E**). Given the central role of TGF-β signaling in liver injury and regeneration^37^, these findings nominated TGF-β as a candidate hepatic niche-derived signal driving progenitor-like reprogramming.

We next asked whether hepatocytes, the dominant parenchymal component of the liver, could provide the ligand source and activate this pathway in tumor cells. TGF-β levels were higher in hepatocyte cultures than in SCLC cells, and SCLC cells exposed to hepatocytes up-regulated TGF-βRII, suggesting increased ligand availability and enhanced tumor-cell responsiveness (**Fig. 2F**). Hepatocyte co-culture activated canonical TGF-β–SMAD signaling in SCLC cells, as shown by increased phospho-SMAD2, and was accompanied by induction of hepatic and progenitor markers (**Fig. 2G**). A similar signaling state was observed *in vivo*, where liver-adapted derivatives showed increased TGF-β and phospho-SMAD2 relative to parental cells (**Fig. 2H**).

We then tested whether TGF-β signaling was sufficient and required for hepatic reprogramming. Exogenous TGF-β induced rapid SMAD2/3 phosphorylation and increased expression of hepatic and progenitor-associated markers, recapitulating the hepatocyte-induced state (**Fig. 2I**). Conversely, pharmacologic inhibition of TGF-β signaling with LY-3200882 suppressed SMAD2 phosphorylation and attenuated induction of reprogramming markers during hepatocyte co-culture (**Fig. 2I**). Genetic disruption of TGF-β receptor signaling similarly impaired phospho-SMAD2 induction and reduced acquisition of the hepatobiliary progenitor-like phenotype, confirming that this response depends on tumor-cell TGF-β pathway responsiveness (**Fig. 2J**).

Finally, we asked whether TGF-β signaling acts on the regulatory elements that define this state. Chromatin immunoprecipitation followed by qPCR after hepatocyte co-culture showed enrichment of phospho-SMAD2 at regulatory regions of HNF4A and SOX9 relative to control loci, with greater enrichment in liver-adapted than parental cells **(Fig. 2K)**. These findings indicate that TGF-β–activated SMAD signaling directly engages regulatory elements associated with hepatic and progenitor identity^38–40^. Together, these data show that hepatocyte-derived TGF-β activates SMAD signaling at hepatic progenitor regulatory loci to promote lineage reprogramming in metastatic SCLC cells.

### Hepatic niche-derived hypoxia potentiates TGF-β signaling to amplify progenitor-like reprogramming in liver metastases

Because hypoxia can cooperate with TGF-β/SMAD signaling^41^, and the liver exhibits pronounced oxygen gradients^36^, we asked whether hypoxia reinforces progenitor-like reprogramming in liver-metastatic SCLC. Consistent with this possibility, VEGFA was among the top inferred ligands in liver-metastatic compared with parental tumor cells (Fig. 2D), and hepatocyte co-culture induced hypoxia-associated programs in tumor cells (fig. S2F). Single-cell transcriptomic analysis further showed that hypoxia pathway activity was elevated in liver-metastatic cells and increased along liver-adapted pseudotime trajectories, whereas parental cells showed comparatively low activity (**Fig. 3A**).

**Fig. 3:**
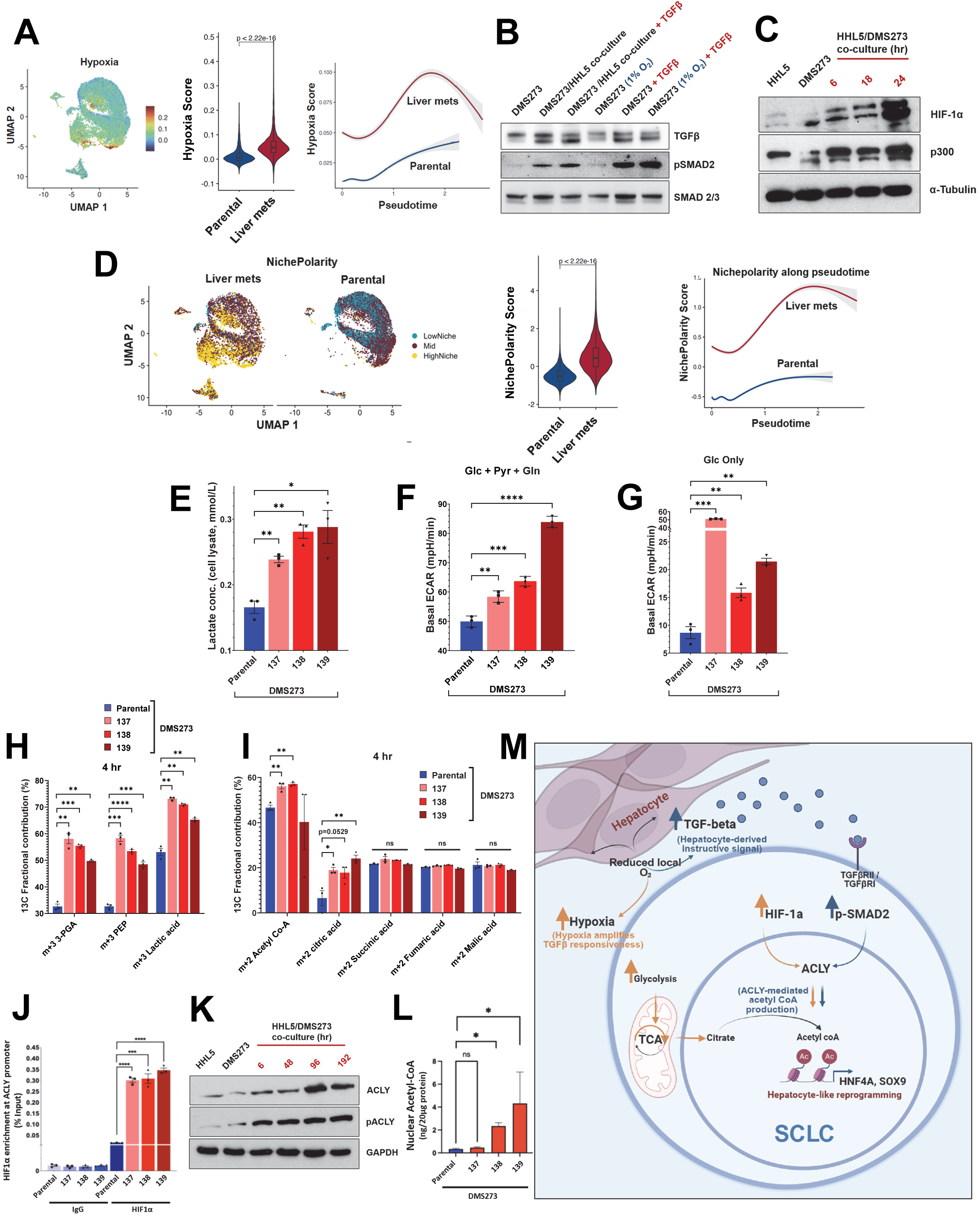
Hepatocyte-driven hypoxia amplifies TGF-β-mediated reprogramming through metabolic–epigenetic coupling. **(A)** UMAP projection, module score distributions, and pseudotime analysis of hypoxia-associated transcriptional programs in parental and liver-metastatic DMS273 cells. **(B)** Immunoblot analysis of DMS273 cells cultured under normoxic or hypoxic conditions with or without TGF-β, probed for phospho-SMAD2 and total SMAD2/3. **(C)** Immunoblot analysis of DMS273 cells co-cultured with HHL5 for the indicated time points, probed for HIF-1α and p300 **(D)** UMAP projection, module score distributions, and pseudotime analysis of NichePolarity scores (integrating z-scored module activities for TGF-β, hypoxia, and HHyP) in parental and liver-metastatic DMS273 cells. **(E)** Intracellular lactate levels in parental and liver-metastatic DMS273 cells (mean ± SEM, n = 3). **(F)** Basal extracellular acidification rate (ECAR) measurements in parental and liver-metastatic DMS273 cells under nutrient-replete, non-limiting conditions (mean ± SEM, n = 3). **(G)** Basal ECAR under glucose-only conditions showing sustained, glucose-dependent glycolytic flux in liver-metastatic cells relative to parental cells (mean ± SEM, n = 3). **(H)** ¹³C-glucose tracing showing increased incorporation of labeled carbon into glycolytic intermediates in liver-metastatic SCLC cells (mean ± SEM, n = 3). **(I)** ¹³C tracing of TCA-associated metabolites showing selective enrichment of labeled acetyl-CoA and citrate, with minimal changes in downstream intermediates (mean ± SEM, n = 3) **(J)** ChIP-qPCR analysis of HIF-1α occupancy at the *ACLY* promoter in parental and liver-metastatic DMS273 cells (mean ± SEM, n = 3). **(K)** Time-course immunoblot analysis of DMS273 cells during hepatocyte co-culture, probed for ACLY and phospho-ACLY. **(L)** Steady-state metabolite analysis of acetyl-CoA levels in nuclear fractions of parental and liver-metastatic DMS273 cells (mean ± SEM, n = 3). **(M)** Working model: hepatocytes establish a local hypoxic microenvironment that amplifies TGF-β signaling in SCLC cells. Hypoxia and TGF-β converge on a HIF-1α–ACLY axis, linking glucose metabolism to acetyl-CoA–dependent epigenetic remodeling and promoting progenitor-like reprogramming. Numbers 137, 138, 139 denote independent liver-metastatic derivatives obtained from separate animals. **Abbreviations:** NichePolarity, Composite score defined as the mean of z-scored TGF-β, hypoxia, and hepatobiliary progenitor (HHyP) module activities; ECAR, Extracellular acidification rate (ECAR); TCA, tricarboxylic acid cycle; Glc, glucose; Pyr, pyruvate; Gln, glutamine. Statistical significance: Two-sided Wilcoxon rank-sum tests or unpaired two-tailed *t*-test; *P* < 0.05 (*), < 0.01 (**), < 0.001 (***), < 0.0001 (****); ns = not significant.

We next tested whether hypoxia modulates TGF-β–SMAD signaling. DMS273 cells were cultured under normoxic or hypoxic conditions, with or without TGF-β stimulation (**Fig. 3B**). Hypoxia modestly increased basal SMAD2 phosphorylation and potentiated the response to TGF-β, with maximal phospho-SMAD2 levels observed under combined hypoxia and TGF-β stimulation. Total SMAD2/3 levels remained unchanged, indicating altered pathway activation rather than changes in SMAD abundance. Consistent with a hypoxia-primed state, SCLC cells in hepatocyte co-culture showed progressive induction of HIF-1α (**Fig. 3C**). Liver-adapted DMS273 cells isolated from *in vivo* metastases similarly exhibited increased HIF-1α protein levels (**fig. S4A**). Chromatin fractionation demonstrated increased nuclear accumulation of HIF-1α in liver-metastatic cells, indicating activation of transcriptionally competent HIF signaling (**fig. S4B**). This was accompanied by up-regulation of the HIF co-activator p300/EP300 (**Fig. 3C, fig. S4A**), a histone acetyltransferase linked to enhancer activation and hepatobiliary differentiation^42,43^.

The enrichment of VEGFA and hypoxia programs in liver-metastatic and hepatocyte co-cultured tumor cells suggested that hepatocytes may contribute to a hypoxic niche that promotes reprogramming. Real-time metabolic profiling showed that human hepatocytes consumed oxygen at higher rates than either parental or liver-adapted SCLC cells, which did not differ significantly from each other under non-limiting conditions (**fig. S4C, D**). Consistent with enhanced oxidative activity, hepatocytes displayed a higher MitoTracker Red-to-Green fluorescence ratio than SCLC cells, reflecting increased mitochondrial membrane potential relative to mitochondrial mass (**fig. S4E**).

To define the ultrastructural basis of this oxidative phenotype, we performed focused ion beam scanning electron microscopy with three-dimensional mitochondrial segmentation and morphometric analysis^44^. Hepatocytes contained highly fused, branched mitochondrial networks with densely packed cristae, features characteristic of high respiratory capacity and sustained oxidative metabolism. By contrast, SCLC mitochondria were smaller and more uniform, with shorter, less densely packed cristae distributed more diffusely throughout the cytoplasm (**fig. S4F**). These data support a model in which metabolically active hepatic parenchyma contributes to local oxygen depletion and hypoxia-associated signaling in metastatic tumor cells. Because hypoxia occurs across many tumor contexts, it is unlikely to explain liver-specific reprogramming on its own. Rather, our data support a cooperative model in which hepatocyte-derived TGF-β provides liver-niche instruction, while local oxygen stress amplifies SMAD signaling and enables HIF-1α–ACLY-dependent metabolic–epigenetic remodeling.

This convergence of hepatobiliary progenitor programs, TGF-β signaling, and hypoxia suggested that liver adaptation reflects a coordinated niche-response state rather than independent pathway activation. To quantify this state, we derived a composite NichePolarity score by integrating z-scored module activities for the fetal HHyP program, TGF-β signaling, and hypoxia. NichePolarity scores were significantly higher in liver-metastatic cells than in parental cells, and increased progressively along liver-metastatic pseudotime trajectories, with minimal change across parental trajectories (**Fig. 3D**). In patient-derived single-cell datasets, HHyP-high tumor cells (fig. S1F) showed significantly greater TGF-β and hypoxia pathway activity than HHyP-low cells (**fig. S4G**), supporting conservation of this coordinated niche-response state in human liver metastases.

### Hypoxia-driven metabolic rewiring fuels acetyl-CoA–dependent epigenetic remodeling in liver-metastasis

Because HIF-1α regulates glucose uptake and glycolytic flux^45^, we asked whether liver adaptation in SCLC was accompanied by metabolic rewiring. Liver-metastatic DMS273 derivatives showed increased intracellular and extracellular lactate relative to parental cells (**Fig. 3E, fig. S4H**), consistent with enhanced glycolytic activity. Liver-adapted RA22 derivatives similarly exhibited increased uptake of the fluorescent glucose analog 2-NBDG and higher intracellular lactate levels compared with parental cells (**fig. S4I, J**).

Real-time metabolic profiling further supported this shift. Liver-adapted SCLC cells displayed higher extracellular acidification rates than parental cells under nutrient-replete conditions (**Fig. 3F, fig. S4K**), without a corresponding increase in oxygen consumption (**fig. S4L**). Increased extracellular acidification persisted in glucose-only medium (**Fig. 3G, fig. S4M**), indicating sustained glucose-dependent glycolytic flux in the liver-adapted state.

To define carbon flow through this pathway, we performed tracing with uniformly labeled ^13^C-glucose (**fig. S4N**). Liver-adapted SCLC cells showed increased ^13^C incorporation into glycolytic intermediates, including 3-phosphoglycerate and phosphoenolpyruvate, and into lactate relative to parental cells (**Fig. 3H**), consistent with enhanced glucose-derived lactate production. Tracing of tricarboxylic acid cycle-associated metabolites revealed selective enrichment of labeled acetyl-CoA and citrate, whereas downstream intermediates, including succinate, fumarate, and malate, were largely unchanged (**Fig. 3I**). Thus, despite HIF-1α–associated constraints on pyruvate dehydrogenase activity^46,47^, liver-adapted cells retained sufficient glucose-derived mitochondrial carbon flux to generate acetyl-CoA and citrate. These data support a hybrid metabolic state in which increased glycolysis coexists with partial glucose oxidation, enabling diversion of carbon toward anabolic pathways and chromatin-associated acetylation. Consistent with this model, lipid accumulation was increased in liver compared with brain metastases in the RA22 model (**fig. S4O**).

Because acetyl-CoA is the required acetyl donor for histone acetylation, we next asked whether liver-associated metabolic rewiring fuels chromatin modification. ^13^C-glucose tracing followed by quantitative mass spectrometry of histone H3 peptides showed increased incorporation of glucose-derived acetyl groups into histones in liver-adapted cells over time, including singly acetylated peptides and the doubly acetylated H3K18ac-K23ac peptide (**fig. S4P, Q**). These data provide direct evidence that glucose-derived carbon contributes to histone acetylation in liver-metastatic SCLC cells.

To probe the link between glucose metabolism and acetyl-CoA–dependent chromatin remodeling, we examined ATP-citrate lyase (ACLY), which converts citrate to acetyl-CoA. ChIP-qPCR demonstrated enrichment of HIF-1α at the *ACLY* promoter in liver-adapted cells, whereas non-target regions and IgG controls showed minimal signal **(Fig. 3J)**. Hepatocyte co-culture reproduced this effect, with increased HIF-1α occupancy at the ACLY promoter compared with monoculture **(fig. S4R)**. Time-course analysis further revealed sequential induction of HIF-1α followed by phospho-ACLY during co-culture **(Fig. 3K)**, consistent with hypoxia-dependent activation of this metabolic node. In agreement with these findings, steady-state metabolite measurements showed increased acetyl-CoA in both cytoplasmic and nuclear fractions of liver-adapted cells **(Fig. 3L, fig. S4S)**, indicating enhanced substrate availability for histone acetylation.

Together, these findings support a model in which the hepatic niche promotes hypoxia-associated signaling that potentiates TGF-β activity and activates a HIF-1α–ACLY metabolic axis in liver-metastatic SCLC cells. This axis couples glucose metabolism to acetyl-CoA–dependent histone acetylation, providing a metabolic–epigenetic mechanism for liver progenitor-like reprogramming (**Fig. 3M**).

### Epigenetic remodeling via acetylation mediates hepatic progenitor-like reprogramming

Because hepatocyte-derived TGF-β and hypoxia converge on the ACLY axis to increase acetyl-CoA availability, we asked whether hepatic niche signals drive coordinated changes in chromatin accessibility and histone acetylation during liver-metastatic reprogramming **(Fig. 4A)**.

**Fig. 4:**
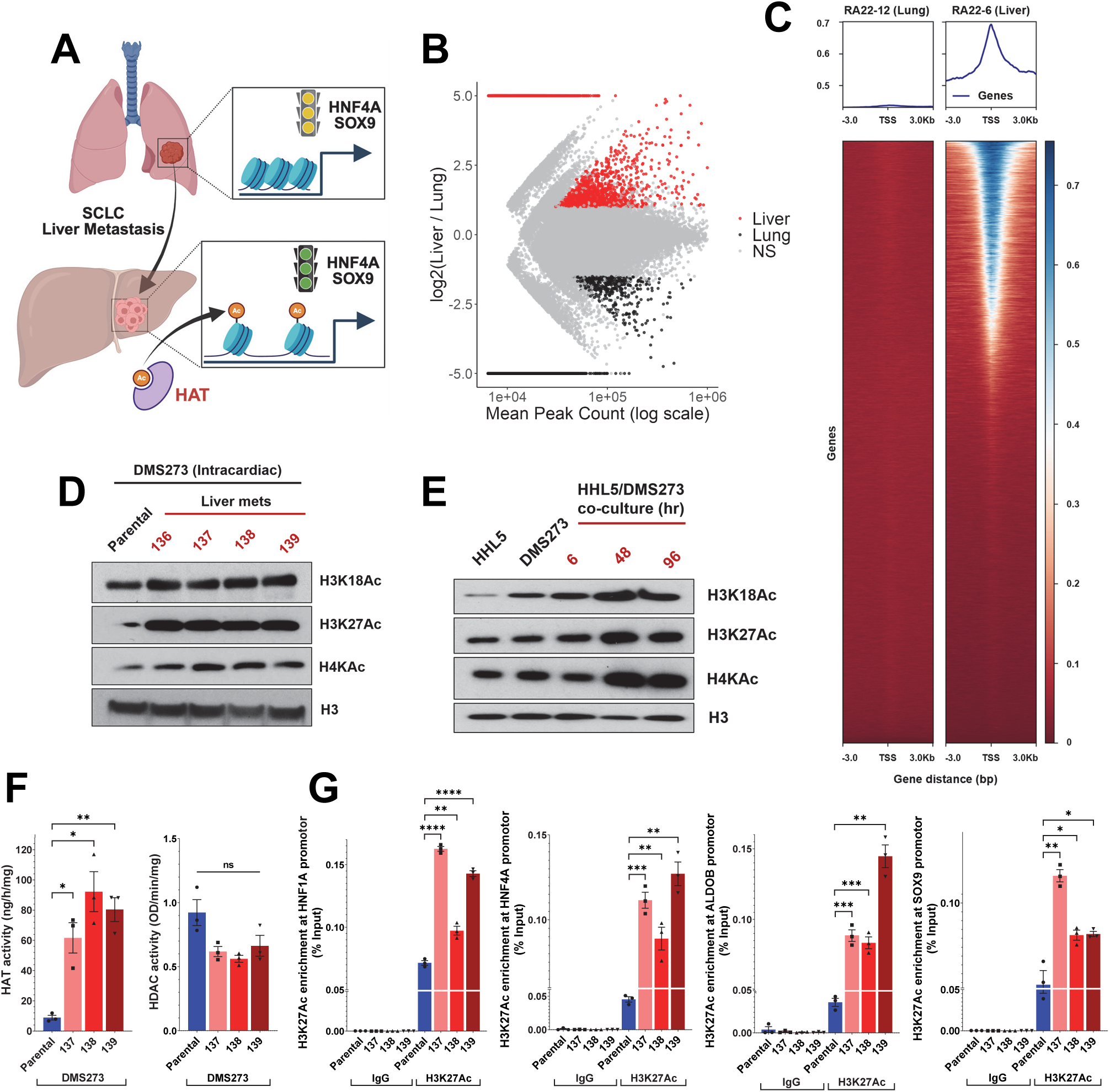
Epigenetic remodeling via acetylation mediates hepatic progenitor-like reprogramming. **(A)** Schematic illustrating chromatin remodeling-driven hepatic progenitor-like reprogramming in SCLC liver metastases. **(B)** ATAC-seq analysis comparing chromatin accessibility in liver and lung metastases from rapid-autopsy SCLC tumors [n = 20 tumors from 11 patients; liver metastases, n = 12; lung lesions, n = 8]. Red, liver-enriched peaks; blue, lung-enriched peaks; gray, not significant. **(C)** Heatmap and average signal profiles of chromatin accessibility centered on transcription start sites in matched liver and lung metastases from the same patient (RA#22). **(D, E)** Immunoblot analysis of acid-extracted histones from parental and liver-metastatic DMS273 cells in the intracardiac model (D), and from DMS273 cells cultured alone or with hepatocytes (E), probed for H3K18ac, H3K27ac, H4Kac, and total histone H3. **(F)** Histone acetyltransferase (HAT) and histone deacetylase (HDAC) activity assays in parental and liver-metastatic DMS273 cells (mean ± SEM, n = 3). **(G)** ChIP-qPCR analysis of H3K27ac enrichment at regulatory regions of *HNF1A*, *HNF4A*, *ALDOB*, and *SOX9* in parental and liver-metastatic DMS273 cells (mean ± SEM, n = 3). Numbers 137, 138, 139 denote independent liver-metastatic derivatives obtained from separate animals. For the DMS273 model (Fig. 4D-F), the parental cells correspond to GFP-labeled DMS273 cells originally isolated from malignant pleural effusion. **Abbreviations:** ATAC-seq, assay for transposase-accessible chromatin using sequencing; HAT, histone acetyltransferase; HDAC, histone deacetylase; TSS, transcription start site; ChIP-qPCR, chromatin immunoprecipitation followed by quantitative PCR. Statistical significance: One-way ANOVA with post hoc Tukey test or unpaired two-tailed *t*-test; *P* < 0.05 (*), < 0.01 (**), < 0.001 (***), < 0.0001 (****); ns = not significant.

We first performed ATAC-seq profiling of SCLC tumors obtained through rapid autopsy, including 20 tumors from 11 patients: 12 liver metastases and 8 lung lesions (**table S6**). Principal component analysis of differentially accessible regions separated liver metastases from lung lesions (**Fig. 4A**). Lung lesions formed a relatively tight cluster, consistent with a more homogeneous accessibility state, whereas liver metastases showed broader dispersion, suggesting increased chromatin-state heterogeneity within the hepatic metastatic niche.

Differential accessibility analysis revealed a predominance of regions with increased accessibility in liver metastases relative to lung tumors, with liver-enriched accessible peaks outnumbering lung-enriched peaks by more than 59-fold (log2FC > 1, adjusted P < 0.05; **Fig. 4B**). Genome-wide heatmaps and metagene plots further confirmed a pronounced increase in accessibility around transcription start sites in liver metastases (**Fig. 4C**). These findings are consistent with prior reports of increased chromatin accessibility in metastatic SCLC in mouse models^48^.

To define the basis of this epigenetic shift, we profiled histone acetylation in matched parental and liver-adapted SCLC cells. Liver-adapted DMS273 cells showed pronounced increases in H3K18ac, H3K27ac, and H4Kac compared with parental cells (**Fig. 4D**). Similar acetylation gains were observed in independent *in vivo* models, including RA22 #6 intracardiac and RA22 #4 intrasplenic liver metastasis models (**fig. S5A, B**). In hepatocyte co-culture, SCLC cells exhibited a time-dependent rise in histone acetylation within 6 hours (**Fig. 4E**), implicating hepatocyte-derived signals as rapid drivers of this epigenetic response. Consistent with these findings, liver-adapted SCLC cells showed increased histone acetyltransferase activity and reduced histone deacetylase activity relative to parental cells (**Fig. 4F**).

We next asked whether these acetylation changes were linked to activation of the hepatic progenitor-like program. ChIP-qPCR showed enrichment of activating histone marks, including H3K27ac and H3K18ac, at regulatory regions of *HNF1A*, *HNF4A*, *ALDOB*, and *SOX9* in liver-adapted cells (**Fig. 4G, fig. S5C**). Notably, chromatin accessibility at these loci showed little change between parental and metastatic cells, suggesting that these regulatory regions are pre-accessible rather than newly opened during metastatic adaptation. Thus, hepatic niche-induced acetylation appears to activate poised lineage-regulatory loci, consistent with injury and regeneration programs in which progenitor-associated genes are rapidly induced through enhancer acetylation and coactivator recruitment^32^.

### ACLY sustains hepatic reprogramming and metastatic fitness in SCLC liver metastases

Because hepatocyte-derived TGF-β and hypoxia converge on a HIF-1α–ACLY metabolic–epigenetic axis, we next asked whether ACLY is required to sustain hepatic reprogramming and metastatic fitness in SCLC liver metastases.

To test this, we generated ACLY knockdown DMS273 cells (**Fig. 5A**) and examined histone acetylation. ACLY depletion markedly reduced global histone acetylation, including H3K18ac and H3K27ac, without altering total histone H3 levels (**Fig. 5B, fig. S6A**). Consistent with impaired epigenetic activation, ACLY knockdown attenuated hepatocyte-induced reprogramming: control cells robustly induced HNF1A, HNF4A, ALDOB, and SOX9 during co-culture, whereas ACLY-depleted cells showed reduced induction of these markers (**Fig. 5C**).

**Fig 5:**
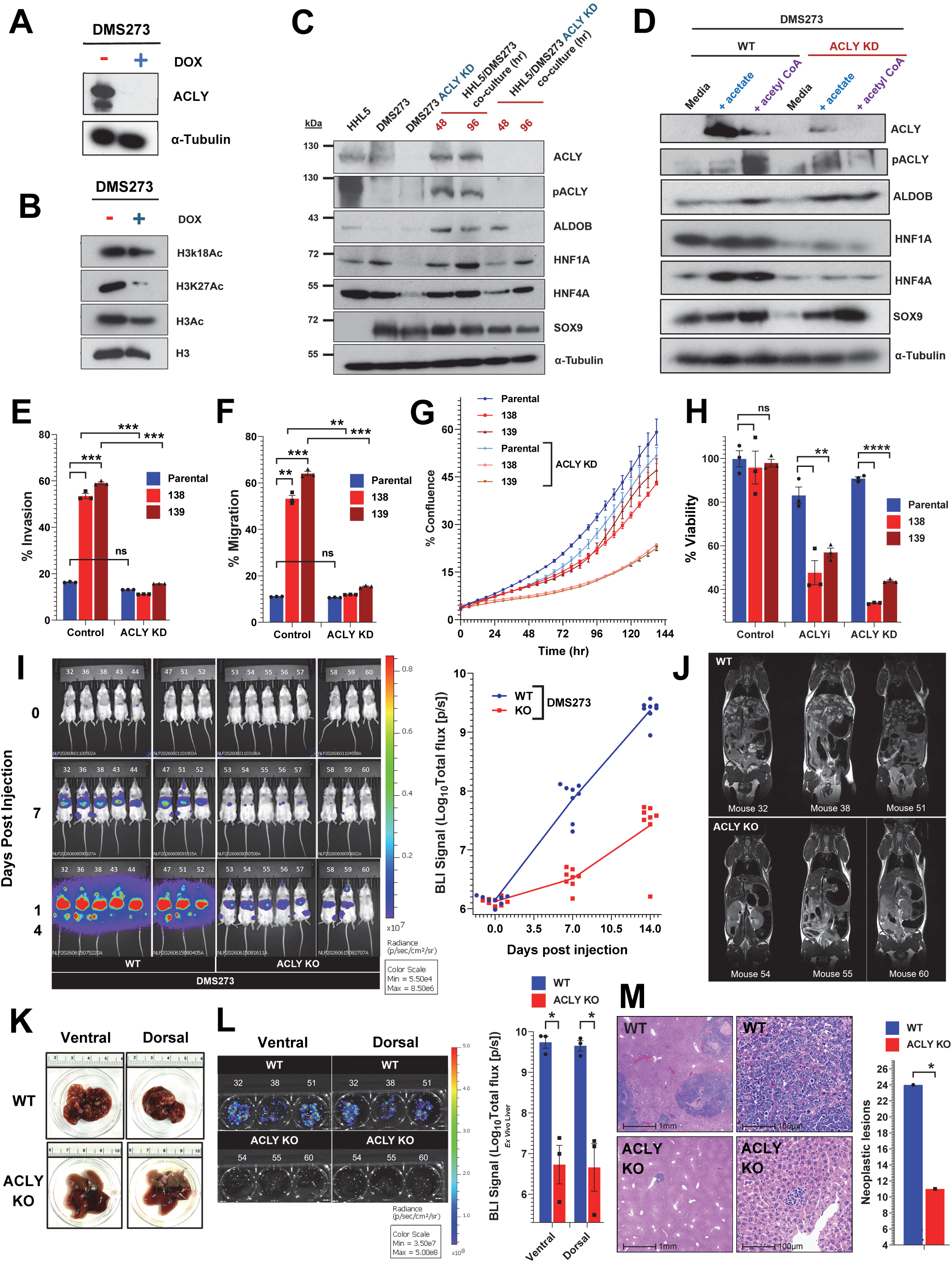
ACLY sustains hepatic reprogramming and fitness in SCLC liver metastases. **(A)** Immunoblot analysis confirming doxycycline-inducible *ACLY* knockdown in DMS273 cells. **(B)** Immunoblot analysis of histone acetylation marks (H3K18ac, H3K27ac, H3Ac, and total H3) in histone lysates from DMS273 cells following *ACLY* knockdown. **(C)** Immunoblot time-course analysis of DMS273 control and ACLY KD cells co-cultured with hepatocytes (HHL5) for the indicated time points (48 and 96 h), probed for ACLY, phospho-ACLY, ALDOB, HNF1A, HNF4A, SOX9, and α-tubulin. **(D)** Immunoblot analysis of DMS273 WT and *ACLY* KD cells following supplementation with sodium acetate or acetyl-CoA, probed for ACLY, phospho-ACLY, ALDOB, HNF1A, HNF4A, SOX9, and α-tubulin. Cells were pre-treated with doxycycline (2 μg/ml, 72 h) prior to metabolite supplementation. **(E, F)** Transwell invasion **(E)** and migration **(F)** assays comparing parental and liver-metastatic DMS273 derivatives following ACLY knockdown. **(G)** Live-cell imaging analysis of proliferative capacity in parental and liver-metastatic DMS273 derivatives following *ACLY* knockdown. **(H)** Effect of pharmacologic ACLY inhibition (ACLYi) and *ACLY* knockdown on viability in parental and liver-metastatic DMS273 derivatives. **(I)** *In vivo* bioluminescence imaging analysis following ultrasound-guided intracardiac injection of DMS273 WT or *ACLY* KO cells into NSG mice. Left, representative BLI images at days 0, 7, and 14 post-injections. Right, quantification of total flux over time. **(J)** Representative MRI images of mice injected with DMS273 WT or ACLY KO cells on day 15 post-injection. **(K)** Representative dorsal and ventral liver images collected on day 15 following intracardiac injection of DMS273 WT or ACLY KO cells. **(L)** Ex vivo liver bioluminescence imaging on day 15 following intracardiac injection of DMS273 WT or ACLY KO cells. Right, quantification of liver-associated BLI signal. **(M)** Hematoxylin and eosin staining of liver sections from mice injected with DMS273 WT or ACLY KO cells. Right, quantification of neoplastic lesions. Numbers 137, 138, 139 denote independent liver-metastatic derivatives obtained from separate animals. **Abbreviations:** SCLC, small-cell lung cancer; ACLY, ATP-citrate lyase; pACLY, phosphorylated ACLY; KD, knockdown; scRNA-seq, single-cell RNA sequencing; H&E, hematoxylin and eosin; IHC, immunohistochemistry; H-score, histological scoring index. Statistics: Paired two-tailed t-test or one-way analysis of variance (ANOVA) with post-hoc Tukey test; P < 0.05 (), < 0.01 (), < 0.001 (), < 0.0001 (****); ns = not significant.

To determine whether this effect reflected reduced acetyl-CoA availability, we supplemented control and ACLY knockdown cells with acetate or acetyl-CoA. Acetyl donor supplementation increased hepatic progenitor marker expression in control cells and partially restored marker induction in ACLY-depleted cells (**Fig. 5D**), accompanied by recovery of H3K18ac and H3K27ac levels (**fig. S6B**). These data indicate that ACLY-dependent acetyl-CoA production is required to maintain histone acetylation and support hepatic progenitor-like reprogramming.

We next asked whether ACLY is activated in human SCLC liver metastases. Immunohistochemical analysis of matched liver and non-liver tumors showed increased ACLY expression in liver lesions, with strongest staining at the tumor-liver interface (**fig. S6C-F**). Single-cell RNA-seq analysis similarly showed an increased fraction of ACLY-high tumor cells in liver metastases compared with lung lesions (**fig. S6G**). These findings support microenvironment-associated activation of ACLY in human liver-metastatic SCLC.

We then examined whether ACLY supports fitness properties of liver-adapted tumor cells. ACLY knockdown markedly reduced invasion in liver-metastatic derivatives but had little effect on parental cells (**Fig. 5E**). Similarly, ACLY depletion impaired migration in liver-adapted cells with minimal effect in parental cells (**Fig. 5F**). Longitudinal live-cell imaging further showed a pronounced reduction in proliferation after ACLY knockdown in liver-metastatic cells, whereas parental cells showed only a modest decrease (**Fig. 5G**). Thus, liver-adapted SCLC cells acquire a selective dependence on ACLY for invasive, migratory, and proliferative capacity.

We next asked whether liver-adapted SCLC cells are selectively vulnerable to pharmacologic ACLY inhibition. ACLY inhibition preferentially reduced viability in liver-metastatic derivatives, with more limited effects in parental cells, recapitulating the direction of the genetic depletion phenotype but with a more modest magnitude of effect (**Fig. 5H**).

Finally, we tested whether ACLY is required for metastatic outgrowth *in vivo*. DMS273 wild-type or *ACLY* knockout cells were introduced into NSG mice by ultrasound-guided intracardiac injection, followed by longitudinal bioluminescence imaging on days 0, 7, and 14. Initial tumor signal was detectable in both groups, indicating that ACLY loss did not prevent early tumor cell seeding. However, wild-type cells expanded progressively over time, whereas *ACLY* knockout cells showed delayed outgrowth and significantly lower total flux by day 14 **(Fig. 5I)**. MRI performed on day 15 further showed reduced tumor burden in the *ACLY* knockout group compared with wild-type controls **(Fig. 5J)**. Endpoint liver analysis confirmed reduced macroscopic metastatic burden in the ACLY knockout group **(Fig. 5K)**. *Ex vivo* liver bioluminescence imaging on day 15 similarly showed reduced tumor signals in the liver following ACLY loss **(Fig. 5M)**, and further histologic assessment showed fewer neoplastic lesions relative to wild-type tumors **(Fig. 5L)**.

Together, these data identify ACLY as a central metabolic-epigenetic node that sustains hepatic reprogramming and metastatic fitness in SCLC liver metastases. By coupling glucose-derived carbon flux to acetyl-CoA-dependent histone acetylation, ACLY enables liver-adapted SCLC cells to maintain a hepatic progenitor-like transcriptional program and support proliferation, invasion, migration, and metastatic expansion.

### NichePolarity defines a liver-specific transcriptional state associated with adverse clinical outcomes in SCLC

Building on our identification of an ACLY-dependent niche-response program in liver-metastatic SCLC, we asked whether this coordinated transcriptional state is associated with clinical outcomes. Because NichePolarity integrates hepatobiliary progenitor-like features with microenvironment-responsive programs, TGF-β signaling and hypoxia, we evaluated its relationship with overall survival in an independent human SCLC cohort^49^.

Higher NichePolarity scores were associated with a trend toward reduced overall survival (log-rank p = 0.064; **Fig. 6A**). This association was driven by the liver metastasis subgroup, where high NichePolarity scores were significantly linked to inferior survival with early separation of survival curves (log-rank p = 0.0068; Fig. 6B), whereas no association was observed in patients without liver metastases (log-rank p = 0.19; Fig. 6C).

**Fig. 6.**
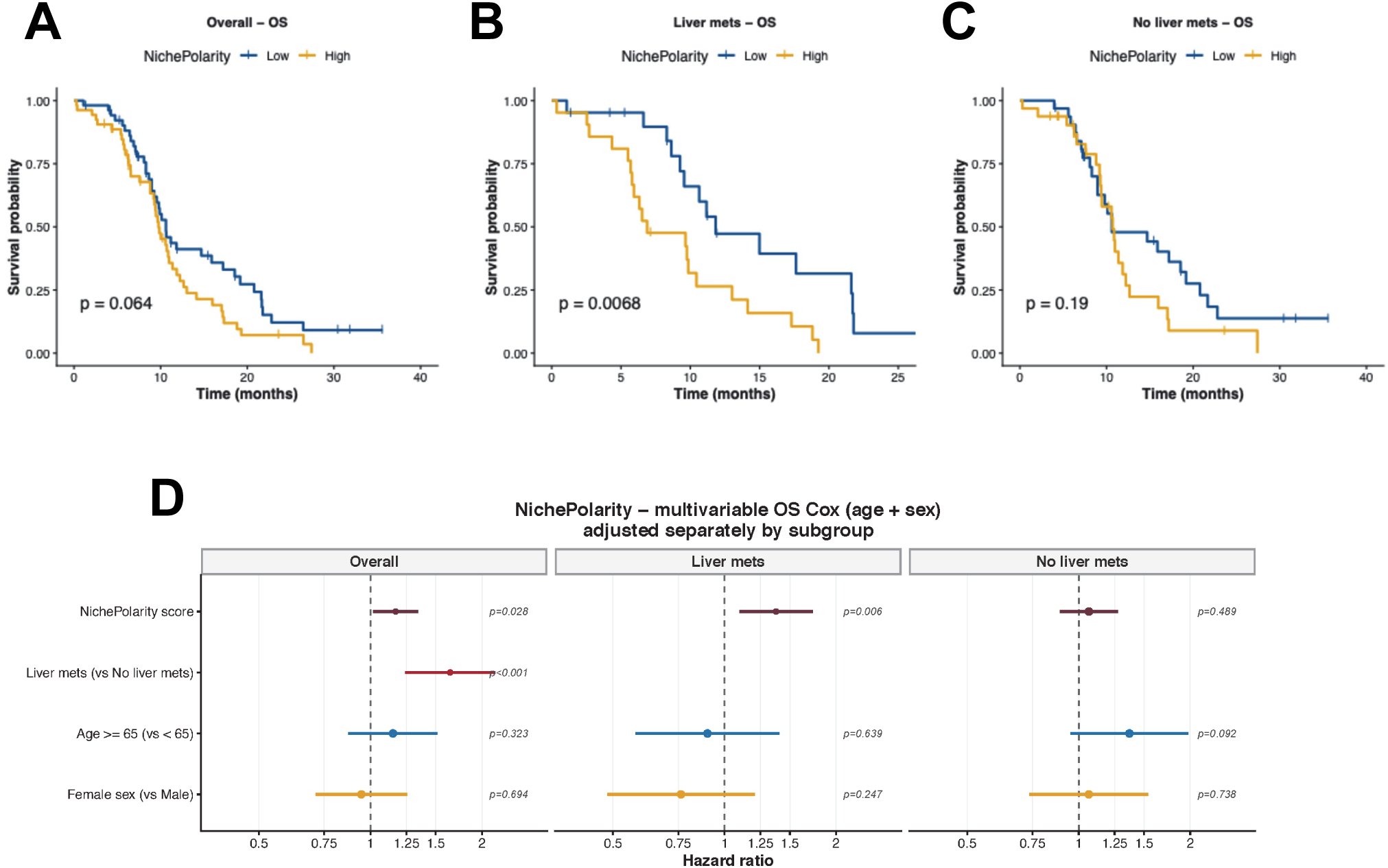
A liver-adapted niche-response state predicts poor survival in patients with SCLC liver metastases. **(A-C)** Kaplan-Meier analyses of overall survival stratified by NichePolarity score (low vs. high) in the Nabet et al overall cohort (A), patients with liver metastases (B), and patients without liver metastases (C). P values were calculated by log-rank test. **(D)** Multivariable Cox proportional hazards analysis of overall survival in the overall cohort and in liver metastasis-defined subgroups, adjusted for age and sex. Hazard ratios are shown with 95% confidence intervals. NichePolarity was modeled as a continuous variable per standard deviation increase.

In multivariable Cox regression adjusting for age, sex, and liver metastasis status, NichePolarity remained an independent predictor of overall survival (HR per SD increase = 1.11, p = 0.028; Fig. 6D), with a stronger effect in the liver metastasis subgroup (HR = 1.43, p = 0.006). No association was observed in patients without liver metastases (p = 0.489). Liver metastasis status remained a significant adverse prognostic factor in the overall cohort (p < 0.001), but NichePolarity contributed additional, non-redundant prognostic information. Age and sex were not significantly associated with overall survival across the overall cohort or within liver and non-liver metastasis subgroups (age p = 0.323, 0.639, 0.092; sex p = 0.694, 0.247, 0.738).

Together, these findings identify NichePolarity as a liver-specific transcriptional state associated with adverse clinical outcomes in SCLC. In the context of our mechanistic framework, this state reflects coordinated activation of TGF-β signaling, hypoxia, and metabolic–epigenetic reprogramming, linking hepatic niche engagement to both tumor cell adaptation and patient survival.

## Discussion

Organ-specific adaptation is a central but incompletely understood dimension of metastasis^52–54^. Our findings show that the liver microenvironment actively reshapes metastatic cell identity by coupling paracrine signaling, oxygen stress, metabolic rewiring, and chromatin remodeling. In SCLC, hepatic colonization induced a reversible hepatobiliary progenitor-like state marked by HNF4A and SOX9. In human SCLC liver metastases, this state was spatially enriched at the tumor–liver interface, could be captured transcriptionally, and was associated with inferior survival among patients with liver metastases. Mechanistically, hepatic niche cues converged through a TGF-β/hypoxia–HIF-1α–ACLY–acetyl-CoA axis to remodel chromatin at hepatic progenitor loci. This state mirrors regenerative dedifferentiation programs activated during liver repair, in which mature hepatocytes can transiently acquire fetal-like progenitor features^31–34,50,51^. We propose that metastatic cancer cells exploit these endogenous hepatic repair circuits, co-opting local tissue cues to reprogram their transcriptional identity. More broadly, hepatobiliary reprogramming in liver metastases illustrates a wider principle: distinct metastatic organs can impose lineage-adaptive states on disseminated cancer cells, analogous to neuronal-like programs reported in brain metastases^50^.

These findings have several implications. First, they reframe liver metastasis as an active process of lineage remodeling driven by hepatic tissue cues. Second, they identify metabolic–epigenetic coupling as a mechanism by which organ-specific signals are converted into cell-state plasticity: TGF-β and hypoxia converge on ACLY-dependent acetyl-CoA production to promote histone acetylation at hepatic progenitor loci. Third, they define ACLY as a functional metabolic–epigenetic node that sustains the liver-adapted state. Finally, they suggest that transcriptional measures of hepatic niche engagement may help identify liver-metastatic tumors most dependent on this adaptive program.

The hepatobiliary progenitor-like program was enriched but not exclusive to liver metastases. In single-cell analyses, fetal HHyP-high tumor cells were approximately twice as frequent in liver lesions as in lung lesions (12.5% versus 6.3%), suggesting that SCLC cells retain baseline lineage plasticity but expansion of this state is favored by the hepatic microenvironment. This interpretation is consistent with recent work in pancreatic cancer showing that malignant cells at the primary site can exhibit transcriptional features of the parenchyma of their ultimate metastatic site^51^. Further, liver reprogramming is a minority, state-specific phenotype rather than a uniform property of all metastatic cells. Nevertheless, its biological relevance is supported by functional data showing that ACLY-dependent liver-adapted cells exhibit increased proliferation, migration, invasion, and metastatic outgrowth, suggesting that even minority niche-adapted populations can disproportionately promote fitness within permissive microenvironments.

Several limitations should be considered. Our experimental metastasis models enabled focused analysis of post-colonization liver adaptation but bypass earlier steps of the metastatic cascade, including invasion, intravasation, and seeding. Although our data implicate hepatocytes as key sources of TGF-β and hypoxia-associated metabolic stress, the liver niche is multicellular, and stellate cells, sinusoidal endothelial cells, Kupffer cells, and recruited immune cells may also shape this state. Additional metabolic and epigenetic mechanisms may contribute to liver-adapted chromatin remodeling. Finally, although ACLY dependency identifies a potential therapeutic vulnerability, clinical translation will require validation in additional liver-metastatic models and assessment of therapeutic windows.

Together, our findings define the liver as an instructive metastatic niche that drives tumor-cell lineage reprogramming through coordinated microenvironmental signaling and metabolic–epigenetic remodeling. By linking hepatic TGF-β and hypoxia to ACLY-dependent acetyl-CoA production, histone acetylation, and activation of hepatic progenitor programs, this work provides a mechanistic framework for organ-specific plasticity in metastasis and identifies liver-adapted metabolic–epigenetic dependencies as potential therapeutic vulnerabilities.

## Supporting information

Supplementary Figures S1 through S6

Supplementary Tables S1 through S6

## Acknowledgements

This research was supported by the Intramural Research Program NCI (ZIA BC 011793) of the National Institutes of Health (NIH). The contributions of the NIH author(s) are considered Works of the United States Government. The findings and conclusions presented in this paper are those of the author(s) and do not necessarily reflect the views of the NIH or the U.S. Department of Health and Human Services. Support from CCR Single Cell Analysis Facility was funded by FNLCR Contract 75N91019D0024. This work utilized the computational resources of the NIH HPC Biowulf cluster. (http://hpc.nih.gov)

## Supplementary Figure Legends

**Fig. S1: SCLC liver metastases acquire liver-associated transcriptional features**

**(A)** Clinical characteristics of patients with SCLC liver metastases compared with non-liver metastatic sites, stratified by age at diagnosis, sex, disease stage, and platinum response.

**(B)** Gene set enrichment analysis of liver-associated transcriptional programs across bulk RNA-seq, patient-matched rapid-autopsy, and proteomic datasets.

**(C)** Oncoprint representation of recurrent mutations and copy-number alterations in liver and non-liver metastases.

**(D)** UMAP visualization of integrated single-cell RNA-seq data showing RB1-loss signature activity across major cellular populations. Violin plots indicate RB1-loss signature scores across annotated cell types.

**(E)** UMAP visualization of adult hepatocyte signature activity across integrated single-cell RNA-seq datasets. Violin plots indicate adult hepatocyte signature scores across annotated cell types.

**(F)** Distribution of fetal HHyP signature scores in tumor cells derived from lung and liver metastases. Red dashed line indicates the 90th-percentile threshold used to define HHyP-high cells.

**(G)** Representative ALDOB immunohistochemistry in rapid autopsy-derived matched liver metastases (RA-25, RA-30, RA-27) and non-liver metastatic sites (lymph node, lung, adrenal gland, pancreas, and pericardial mass). Lower panels, quantification of ALDOB staining across matched metastatic sites. Data are mean ± s.e.m.; statistical analysis by two-tailed paired t-test.

**(H)** Representative ALDOB immunohistochemistry from a rapid-autopsy liver metastasis showing expression at the tumor-liver interface. Insets show higher-magnification views of the indicated region.

**Fig. S2: Experimental models recapitulate liver progenitor-like reprogramming. A,**

**(A)** Flow cytometry analysis of GFP-labeled DMS273 cells and RFP-labeled HHL5 hepatocytes at the indicated co-culture time points, showing separation of tumor and hepatocyte populations.

**(B)** Schematic of indirect transwell co-culture and sorting workflow (left). Immunoblot analysis of DMS273 cells following indirect co-culture with HHL5 hepatocytes for the indicated time points, probing markers selected to capture hepatic lineage specification and progenitor-like reprogramming: HNF4A, a hepatocyte-lineage transcription factor; SOX9, a hepatobiliary progenitor/cholangiocyte-associated factor; and ALDOB, a liver-enriched metabolic enzyme. α-tubulin is shown as a loading control.

**(C)** Immunoblot analysis of H69 cells following indirect co-culture with HHL5 hepatocytes, probed for HNF1A, HNF4A, ALDOB, SOX9, and α-tubulin.

**(D)** Immunoblot time-course analysis of DMS273 cells removed from hepatocyte co-culture and cultured independently for the indicated durations, probed for HNF1A, HNF4A, ALDOB, SOX9, and α-tubulin.

**(E)** RNA-seq analysis of FACS-isolated GFP⁺ DMS273 cells following 6 h hepatocyte co-culture, showing early induction of selected progenitor and injury-response genes, including SPP1, CD44, SOX9, and SOX4, relative to monoculture controls.

**(F)** GSEA of RNA-seq from hepatocyte co-cultured DMS273 cells, showing enrichment of fetal HHyP, TGF-β, hypoxia, Wnt, and VEGF signaling programs, consistent with activation of hepatic injury-response and progenitor-like reprogramming.

**(G)** Immunoblot analysis of HCT116 cells following co-culture with HHL5 hepatocytes, probed for HNF1A, HNF4A, ALDOB, SOX9, and α-tubulin.

**(H)** Principal component analysis of transcriptomic (left) and proteomic (right) profiles from parental (P0), liver-metastatic (Liver.P1), and brain-metastatic (Brain.P1) tumors.

**(I)** Gene set enrichment analysis of the fetal HHyP signature comparing liver-metastatic (P1_Liver) and parental liver-derived (P0_Liver) tumors.

**(J)** Gene set enrichment analysis of the fetal HHyP signature comparing liver-metastatic (P1_Liver) and brain-metastatic (P1_Brain) tumors.

**Fig. S3: Hepatic microenvironment induces conserved progenitor-like reprogramming in liver metastases.**

**(A)** Flow cytometry analysis of GFP⁺ parental and liver-metastatic DMS273 tumors isolated following intracardiac injection into NSG mice.

**(B)** Principal component analysis of bulk RNA-seq profiles from parental and liver-metastatic DMS273 tumors.

**(C)** Immunoblot analysis of RA22#4 patient-derived SCLC cells following intrasplenic injection and liver colonization, probed for HNF1A, HNF4A, ALDOB, SOX9, and α-tubulin. Schematic illustrates the experimental workflow for generation of liver metastases.

**Fig. S4: Metabolic reprogramming in SCLC**

**(A)** Immunoblot analysis of DMS273 parental and liver-metastatic derivatives probed for HIF-1α, p300, ACLY, phospho-ACLY (Ser455), and α-tubulin.

**(B)** Immunoblot analysis of nuclear extracts from parental and liver-metastatic DMS273 cells probed for chromatin-bound HIF-1α. α-tubulin was used as control.

**(C)** Representative basal oxygen consumption rate (OCR) traces in HHL5 hepatocytes, parental DMS273 cells, and liver-metastatic derivatives (mean ± SEM, n = 3 per cell type).

**(D)** Quantification of basal OCR measurements (mean ± SEM, n = 3).

**(E)** Quantification of MitoTracker Red-to-Green fluorescence ratios in HHL5 hepatocytes and DMS273 cells.

**(F)** Focused ion beam scanning electron microscopy (FIB-SEM) and 3D mitochondrial segmentation analysis of HHL5 hepatocytes and DMS273 cells. Representative images show mitochondrial ultrastructure and cristae organization.

**(G)** Analysis of patient tumor scRNA-seq data (Fig. 1 D-G) comparing TGF-β and hypoxia in liver metastatic cells with HHyP-high (above 90^th^ percentile) vs. HHyP-low activity (Top panel) and density distribution of NichePolarity scores in parental and liver-metastatic DMS273 cells (bottom panel).

**(H)** Extracellular lactate measurements in parental and liver-metastatic DMS273 cells (mean ± SEM, n = 3).

**(I)** Quantification of 2-NBDG uptake in parental and liver-adapted RA22 derivatives (mean ± SEM, n = 3).

**(J)** Intracellular lactate measurements in parental and liver-adapted RA22 derivatives (mean ± SEM, n = 3).

**(K)** Representative basal extracellular acidification rate (ECAR) traces in parental and liver-metastatic DMS273 cells (mean ± SEM, n = 3 per cell type).

**(L)** Quantification of basal OCR measurements under non-limiting conditions (mean ± SEM, n = 3).

**(M)** Real-time ECAR measurements in parental and liver-metastatic DMS273 cells cultured in glucose-only medium (mean ± SEM, n = 3 per cell type).

**(N)** Schematic of uniformly labeled ^13^C-glucose tracing illustrating carbon flow through glycolysis and downstream metabolic intermediates.

**(O)** Representative H&E and lipid staining of parental RA22 tumor cells, brain metastases, hepatocytes, and liver metastases. Right, quantification of lipid droplet accumulation across samples, showing increased lipid accumulation in liver metastases relative to parental and brain-metastatic tumors.

**(P)** ^13^C-glucose tracing coupled with quantitative histone mass spectrometry showing incorporation of glucose-derived acetyl groups into doubly acetylated H3K18ac-K23ac peptides at 4 h and 24 h in parental and liver-metastatic DMS273 cells.

**(Q)** Quantification of ^13^C-acetyl incorporation into singly acetylated histone H3 peptides (H3K18ac or H3K23ac) in parental and liver-metastatic DMS273 cells at 24 h.

**(R)** ChIP-qPCR analysis of HIF-1α occupancy at the ACLY promoter in DMS273 cells cultured alone or in hepatocyte co-culture. IgG and non-target genomic regions were used as controls (mean ± SEM, n = 3).

**(S)** Quantification of acetyl-CoA levels in cytoplasmic fraction of parental and liver-metastatic DMS273 cells (mean ± SEM, n = 3).

**Fig. S5: Liver metastatic adaptation is associated with increased histone acetylation at hepatic progenitor loci**

**(A, B)** Immunoblot analysis of acid-extracted histones from parental and liver-metastatic RA22#6 intracardiac **(A)** and RA22#4 intrasplenic derivatives **(B)**, probed for H3K18ac, H3K27ac, H4Kac, and total histone H3.

**(C)** ChIP-qPCR analysis of H3K18ac enrichment at the promoters of *HNF1A*, *HNF4A*, *ALDOB*, and *SOX9* in parental and liver-metastatic DMS273 cells (mean ± SEM, n = 3).

**Fig. S6: ACLY regulates histone acetylation in liver-adapted SCLC cells.**

**(A)** Immunoblot analysis of histone acetylation marks following ACLY knockdown during hepatocyte co-culture, probed for H3K18ac, H3K27ac, and total histone H3.

**(B)** Immunoblot analysis of histone acetylation following acetate or acetyl-CoA supplementation in ACLY KD cells, probed for H3K18ac, H3K27ac, and total histone H3.

**(C)** Representative H&E and ACLY immunohistochemistry of patient-matched lung and liver metastases from patients with SCLC. Insets show low-magnification views. Scale bars, 100 μm.

**(D)** Quantification of ACLY immunohistochemical staining (H-score) in patient-matched lung and liver metastases from patients with SCLC (n = 10 pairs). Statistical analysis by paired two-tailed t-test.

**(E)** Representative ACLY immunohistochemistry across distinct tumor regions in SCLC liver metastases. Insets show higher-magnification views of the indicated regions.

**(F)** Quantification of ACLY immunohistochemical intensity across tumor core, tumor-liver interface, and adjacent hepatocyte regions. Tumor-liver interface was defined as the region within 100 μm of the tumor boundary.

**(G)** Quantification of ACLY-high tumor cells in human single-cell RNA-seq datasets from lung and liver metastases. ACLY-high cells were defined using a threshold of >1.09 expression units.

## Methods

### Ethics Approval and Consent

Tumor samples were obtained under NIH protocols NCT02146170, *Tissue Procurement and Natural History Study of People With Non-Small Cell Lung Cancer, Small Cell Lung Cancer, Extrapulmonary Small Cell Cancer, Pulmonary Neuroendocrine Tumors, and Thymic Epithelial Tumors*, and NCT01851395, *Inpatient Hospice With Procurement of Tissue on Expiration in Thoracic Malignancies, Bladder Cancer, Ovarian Cancer, Epithelial Cancer, and Patients Treated With an Adoptive Cellular Therapy*. These studies were approved by the NIH Institutional Review Board and the Office of Human Subjects Research Protections at the National Cancer Institute under protocols 14-C-0105 and 13-C-0131. All patients provided written informed consent for tumor sequencing and tissue collection in accordance with the Declaration of Helsinki.

### Human Tissue Collection and Processing

Metastatic tumor tissues were collected at biopsy under NCT02146170 or within two hours post-mortem as part of a rapid autopsy program under NCT01851395. Fresh specimens were fixed in 10% neutral buffered formalin followed by paraffin embedding or preserved in MACS Tissue Storage Solution (Miltenyi Biotec, Cat. No. 130-100-008) on ice for immediate processing. Fresh tissue specimens were enzymatically dissociated using the Tumor Dissociation Kit, human (Miltenyi Biotec, Cat. No. 130-095-929), in combination with the gentleMACS Dissociator and the pre-set 37°C hTDK-1 program. High-viability single-cell suspensions were assessed for quality and used for single-cell RNA sequencing. Parallel aliquots of dissociated cells were cultured in RPMI-1640 medium supplemented with 10% fetal bovine serum, 1X GlutaMAX, and 1% penicillin-streptomycin to generate patient-derived primary cell lines.

### Histology and Immunohistochemistry

Formalin-fixed, paraffin-embedded tissue sections were cut at 4 µm thickness for hematoxylin and eosin staining and immunohistochemistry. Immunohistochemistry was performed using the Bond RX autostainer (Leica Biosystems) with Epitope Retrieval 1, citrate buffer, for 20 minutes. Slides were incubated with primary antibodies against Aldolase B (Invitrogen, PA5-51694; 1:1000, 30 minutes) and ATP-citrate lyase (ACLY; Abcam, ab40793; 1:100, 30 minutes). Detection was carried out using the Bond Polymer Refine Detection Kit (Leica Biosystems, DS9800), according to the manufacturer’s instructions. Negative controls included isotype-matched antibody reagents. After staining, slides were dehydrated through graded ethanol, cleared in xylene, and coverslipped. Hematoxylin and eosin staining was performed using the Tissue-Tek Prisma autostainer.

### Slide Digitization and Image Acquisition and Quantitative Image Analysis

Whole-slide images were acquired at 20x magnification using a Carl Zeiss AxioScan Z1 slide scanner. Digital pathology analysis was performed using HALO image analysis software (v3.4.2986.151; Indica Labs). The Tissue Classifier module, based on a Random Forest algorithm, was trained on pathologist-annotated samples to segment tumor, stroma, necrosis, and background. The Cytonuclear module was used to quantify nuclear and cytoplasmic DAB staining intensity. Immunohistochemistry positivity thresholds were optimized using control tissues. The Tissue Microarray module enabled automated core identification and artifact exclusion for batch processing. Cell segmentation was based on DAPI nuclear staining with the following parameters: nuclear size 9–571 µm², minimum roundness 0.05, cytoplasm radius 5 µm, nuclear contrast threshold 0.5, intensity threshold 0.09, and segmentation aggressiveness 0.2. Quantitative outputs included percent positive cells and H-score. Quantitative data from HALO were exported into GraphPad Prism 10.4.1 for statistical analysis

### Tumor RNA-seq, Whole Exome Sequencing, and ATAC-seq

FFPE or frozen tumor tissue samples were used for both RNA-seq and whole-exome sequencing (WES). For WES, 100 ng of DNA was sheared (∼200 bp; Covaris), and exome enrichment was performed using the SureSelect Clinical Research Exome Kit (Agilent). RNA was extracted using the RNeasy FFPE kit (QIAGEN), and RNA-seq libraries were generated using the TruSeq RNA Exome Library Prep Kit (Illumina). Raw FASTQ files were processed using Trimmomatic to remove adapter sequences and low-quality reads, followed by alignment to the hg19 reference genome using STAR (v2.4.2a). Gene expression levels were quantified using Rsubread featureCounts, and normalized to Transcripts Per Million (TPM) to enable expression comparison across samples. For somatic mutation analysis, sequencing reads were aligned to hg38 using BWA-MEM, followed by duplicate removal and base recalibration with GATK. Strelka2 was used for somatic SNV and INDEL calling, and filtered variants were annotated with VEP to predict functional impact.

For autopsy-derived metastatic tumor tissues, ATAC-seq was performed using the Illumina HiSeq 4000 platform (Novogene). Raw reads were processed using the nf-core/atacseq pipeline (v2.1.2). Quality control was performed with FastQC, and adapters were trimmed using Trim Galore. Reads were aligned to the hg38 reference genome using BWA-MEM. PCR duplicates were marked with Picard and reads mapping to mitochondrial DNA or with ambiguous alignment were removed. Accessible chromatin peaks were identified using MACS2 with ATAC-specific parameters. Consensus peak sets were generated across samples to identify reproducible regulatory regions. Peak matrices were used for downstream visualization, including promoter accessibility heatmaps generated using DeepTools.

To evaluate global differences in chromatin accessibility between liver metastatic and lung tumor samples, principal component analysis (PCA) was performed on a subset of differentially accessible regions. Rather than using all genome-wide accessible peaks, dimensionality reduction was restricted to the most variable features to enhance biological signal and reduce contributions from constitutively accessible regions. Specifically, the top 1% of differentially accessible peaks (ranked by variance, with log2 fold change and adjusted p-value < 0.05) were selected for downstream analysis. Accessibility counts were normalized using variance stabilizing transformation (VST), and the resulting matrix was used as input for PCA. Principal components were computed using standard singular value decomposition as implemented in R. Pairwise similarity between samples was assessed using Pearson correlation coefficients calculated on the same VST-normalized accessibility matrix. Correlation values were visualized as a sample-by-sample similarity matrix to evaluate inter-sample concordance across tumor groups.

### Single-Cell RNA Sequencing

Single-cell suspensions were processed for single-cell RNA-seq gene expression profiling according to the Chromium GEM-X Single Cell 3’ v4 User Guide (10x Genomics, CG000731 Rev B). Cell counts and viability were assessed using acridine orange/propidium iodide fluorescent dyes (LGBD10012) on the Luna-FX7 Automated Cell Counter (Logos Biosystems). Sample concentrations were adjusted as needed by dilution in PBS with 0.04% BSA. Cells were partitioned and barcoded using the Chromium X platform (10x Genomics), targeting 12,000 cells per sample. Sequencing libraries were generated and quantified using the Chromium Connect automated platform (10x Genomics).

Sequencing was performed on an Illumina NovaSeq X Plus instrument using a 100-cycle kit with 10x Genomics-recommended run parameters of 28-10-10-90. The average target read depth was approximately 50,000 reads per cell. Base calling was performed using RTA4, and demultiplexing was conducted using BCL Convert v4.1.23. Data were processed using the 10x Genomics Cell Ranger pipeline v9.0.1 with STAR 2.7.2a. Sequenced reads were aligned to a custom reference using the 10x Genomics-provided human reference, refdata-gex-GRCh38-2024-A, modified to include GFP sequence.

### Single-Cell Transcriptomic Analysis of Human SCLC

Single-cell transcriptomic data from SCLC samples were analyzed using Scanpy (v1.9.3)^52^ in Python. Preprocessed .h5ad files were imported into an AnnData object. After incorporating clinical metadata, only samples annotated as “Small Cell Lung Cancer” from liver or lung tissue sites were retained. Cells with fewer than 100 counts and mitochondrial genes, identified by the prefix “MT-”, were excluded. Expression values were normalized to 10,000 transcripts per cell, followed by natural log transformation using log1p. The resulting log-normalized values were stored as a separate layer.

The top 5,000 highly variable genes were identified per dataset. To mitigate batch effects, Harmony integration^53^ was performed on PCA embeddings derived from log-normalized highly variable genes using sample identity as the batch key, with up to 25 iterations for convergence. Based on the Harmony-corrected principal components, a k-nearest-neighbor graph was constructed, a UMAP embedding was generated for visualization, and unsupervised Leiden clustering was performed at multiple resolutions, including 0.2–1.0.

Clusters were manually annotated by comparing canonical marker genes to identify major cell types, including tumor cells, hepatocytes, myeloid cells, T cells, B cells, endothelial cells, fibroblasts, and undefined cells. Undefined cells were excluded where appropriate. Functional signatures from a .gmt file were scored using the AUCell method through Decoupler on the log-normalized expression matrix. Activity scores for key programs, including RB1-loss^29^ and hepatocyte-related gene sets^54^, were visualized using violin and UMAP plots. Cells exhibiting an RB1-loss score > 0.025 were classified as high-confidence tumor cells and subsetted for focused analysis. Within this tumor subset, marker gene expression and pathway scores were compared between lung and liver metastatic sites using violin plots. The proportion of cells exceeding the 90th percentile expression threshold for specific features was quantified and visualized as bar plots to highlight site-specific adaptations.

### Single-Cell Transcriptomic Analysis

Pathway and lineage activities were quantified at single-cell resolution using AUCell through Decoupler and Seurat AddModuleScore^55–57^. Gene programs included hepatobiliary hybrid progenitor, TGF-β, and hypoxia signatures. Transcription factor activity was inferred using the DoRothEA/VIPER framework^58^.

A composite niche score, termed NichePolarity, also referred to as NicheComposite, was defined as the mean of z-scored TGF-β, hypoxia, and hepatobiliary hybrid progenitor program activities. Cells in the top 20% of NichePolarity values were classified as HighNiche, cells in the bottom 20% as LowNiche, and the remaining cells as Mid. Pseudotime trajectories were analyzed in parental and liver-metastatic DMS273 cells to assess temporal activation of lineage and niche programs. Program dynamics along pseudotime were modeled using generalized additive models with mgcv and k = 8. Program onset was defined as the first pseudotime point at which the fitted curve reached 15% of its dynamic range above baseline. Temporal ordering was summarized by onset and peak pseudotime. Confidence intervals were estimated using bootstrap resampling with 200 iterations.

Ligand-activity inference was performed using NicheNet to identify candidate upstream regulators associated with liver adaptation^59^. Liver-metastatic and parental tumor cells were compared across the full tumor population. Ligand activities were ranked using NicheNet output metrics, including AUPR-corrected scores and AUROC.

### Survival Analysis of the NichePolarity Score (Barzin et al. bulk RNA-seq cohort)

Gene expression data and clinical metadata were obtained from the Barzin et al. bulk RNA-seq dataset comprising 271 SCLC patients^49^. Raw transcripts-per-million values were log_2_-transformed as log_2_(TPM + 1) before scoring. Seven patients with missing LDH values were excluded, leaving a final analytic cohort of 264 patients.

The NichePolarity score was derived as a composite of three transcriptional modules described above: TGF-β signaling, hypoxia, and the hepatobiliary hybrid progenitor program. For each module, single-sample gene set enrichment analysis scores were computed from the log2-transformed bulk RNA-seq matrix using the GSVA package (v2.5.15), with alpha = 0.25 and score normalization enabled. Only genes present in the bulk expression matrix were included, with a minimum of three genes required per module. Module scores were then z-scored independently across samples, and the NichePolarity composite score was calculated as the row-wise mean of the three z-scored module activities.

Associations between NichePolarity score and overall survival and progression-free survival were assessed using Cox proportional hazards regression implemented in the survival package. A multivariable Cox model including liver metastasis status, age category, and sex as covariates was used to assess whether NichePolarity remained an independent predictor after adjustment for key variables of biological relevance. Liver metastasis status was defined using MLIVER; age category was defined as AGE65, comparing patients <65 versus ≥65 years. All continuous scores were standardized to mean 0 and standard deviation 1 before entry into Cox models, so hazard ratios reflect the effect per one standard deviation increase in NichePolarity score.

Analyses were conducted separately in the overall cohort and in subgroups defined by liver metastasis status: liver metastases present, MLIVER = Y, n = 105, and liver metastases absent, MLIVER = N, n = 159. For visualization, patients were stratified into high and low NichePolarity groups. Survival curves were estimated using the Kaplan-Meier method and compared using the log-rank test. Results are displayed as Kaplan-Meier curves with P values from the log-rank test.

### Cell Lines and Culture conditions

The SCLC cell line DMS273 was obtained from MilliporeSigma (Cat. No. 95062830-1VL). Additional cell lines included NCI-H69 (Cat. No. HTB-119), HCT116 (Cat. No. CCL-247EMT), MCF7 (Cat. No. HTB-22), and BEAS-2B/CRL9609 (Cat. No. CRL-3588), all purchased from the American Type Culture Collection. Human immortalized hepatocytes, HHL5, were generously provided by the Arvind H. Patel laboratory^60^.

DMS273, HCT116, MCF7, and HHL5 cells were cultured in Dulbecco’s Modified Eagle Medium supplemented with 10% FBS and 1% penicillin-streptomycin at 37°C in a humidified incubator with 5% CO₂. H69 cells were maintained in RPMI-1640 medium supplemented with 10% FBS, 1X GlutaMAX, and 1% penicillin-streptomycin under the same culture conditions. Patient-derived primary cultures RA22#4, RA22#6, and RA22#12, along with in vivo model-derived liver metastases #400L and #404L and brain metastases #406B and #408B, were generated in-house and cultured in RPMI-1640 medium supplemented with 10% FBS, 1X GlutaMAX, and 1% penicillin-streptomycin at 37°C in 5% CO₂. All cell lines were routinely tested for mycoplasma contamination using the MycoAlert Mycoplasma Detection Kit (Lonza).

### Generation of Metastasis-Derived Cell Lines

Following euthanasia, tumors from liver and brain metastases were harvested and preserved in MACS Tissue Storage Solution (Miltenyi Biotec, Cat. No. 130-100-008) on ice. Tumors were dissociated using the Tumor Dissociation Kit, human (Miltenyi Biotec, Cat. No. 130-095-929), and the gentleMACS Dissociator with the 37°C hTDK-1 protocol. Single-cell suspensions were cultured in RPMI-1640 medium supplemented with 10% FBS and 1% penicillin-streptomycin under standard conditions of 37°C and 5% CO₂ to establish primary cell lines.

### Generation of H2B-GFP and H2B-RFP Expressing Cell Lines

Lentiviral vectors encoding histone H2B fused to fluorescent proteins were used to generate fluorescently labeled cell lines. Lentiviruses were produced by co-transfecting HEK293T cells with the packaging plasmids pCMV-dR8.2 dvpr (Addgene #8455), pCMV-VSV-G (Addgene #8454), and the transfer vectors pHIV-H2B-eGFP (Addgene #91776) or pHIV-H2B-mRFP (Addgene #18982). Transductions were performed using Turbofectin 8.0 (OriGene, TF81001). Viral supernatants were harvested at 48 and 72 hours post-transfection, filtered, pooled, and stored at -80°C.

Target cells, 1-2 million cells per condition, were incubated with polybrene at 2 µg/mL for 10 minutes before infection with 1 mL viral supernatant. After 48 hours, cells were washed with PBS and cultured in DMEM supplemented with 10% FBS. Cells were expanded for five days before fluorescence-activated cell sorting to enrich for GFP- or RFP-positive populations. Stable expression of H2B-GFP or H2B-RFP was confirmed by fluorescence microscopy.

### Hepatocyte-SCLC Co-culture and FACS Sorting

SCLC cell lines H69 and DMS273 were stably labeled with H2B-GFP, while human immortalized hepatocytes HHL5 were labeled with H2B-RFP before co-culture. Cells were plated at a 1:2 ratio of SCLC cells to hepatocytes; however, because of differential growth kinetics, the final cell ratio approached approximately 1:1 at the time of analysis.

Co-culture experiments were conducted for 6 hours, 2 days, or 4 days without media replenishment. For direct co-culture, cells were seeded together in the same well. For indirect co-culture, cells were physically separated using Transwell inserts with 0.4 µm pore size (Costar #7910), allowing exchange of soluble factors without direct contact.

At designated time points, cells were harvested, washed twice with PBS, and pelleted at 5000 rpm for 5 minutes. Mixed cell populations were resuspended in PBS supplemented with 1% FBS and 1 mM EDTA and sorted based on GFP/RFP fluorescence using a BD FACSAria Fusion cell sorter. Sorted DMS273 and H69 SCLC cells were used for whole-cell protein lysate preparation and total RNA isolation using the mirVana RNA Isolation Kit (Invitrogen, AM1914).

### Generation of ACLY Knockdown Constructs and Stable Cell Lines

Short hairpin RNA sequences targeting human ACLY were designed using the Broad Institute GPP Web Portal. Two effective sequences were selected: shRNA-1, 5′-CGTGAGAGCAATTCGAGATTA-3′, and shRNA-2, 5′-GCCTCAAGATACTATACATTT-3′. These sequences were cloned into the doxycycline-inducible lentiviral expression vector pRSIT16-U6Tet-sh-CMV-TetRep-2A-TagRFP-2A-Puro (Cellecta, Cat. #SVSHu6T16-L), according to the manufacturer’s protocol. Cloned plasmids were verified by Sanger sequencing.

Lentiviral particles were generated by co-transfecting HEK293T cells with the shRNA plasmid, pCMV-VSV-G (Addgene #8454), and pCMV-dR8.2 dvpr (Addgene #8455) using Turbofectin 8.0 (OriGene, TF81001). Viral supernatants were harvested at 48 and 72 hours post-transfection, pooled, filtered, and stored at -80°C. Target SCLC cells, 1-2 million per well, were pre-treated with 2 µg/mL polybrene for 10 minutes and transduced with 1 mL viral supernatant. After 48 hours, cells were washed and cultured in fresh DMEM with 10% FBS for two additional days, followed by selection with 2 µg/mL puromycin for five days. RFP-positive cells were subsequently sorted by flow cytometry to enrich for transduced populations. ACLY knockdown efficiency was validated by western blotting.

### Migration Assay

Cell migration was assessed using a 96-well transwell migration assay with 8 µm pore size (Abcam, ab235673). DMS273 parental and liver-metastatic derivatives 138 and 139 were cultured to approximately 80% confluence and serum-starved for 18-24 hours before the assay. Cells, 5 x 10⁴ per condition, were suspended in serum-free medium and seeded into the upper chamber, while the lower chamber contained chemoattractant-containing medium. Following incubation for 48 hours at 37°C in 5% CO₂, non-migratory cells were removed from the upper chamber. Migrated cells were collected from the lower chamber, stained with fluorescent dye, and quantified by fluorescence measurement at Ex/Em = 530/590 nm. Fluorescence values were converted to cell numbers using a standard curve generated by serial dilution of known cell densities. Migration was expressed as the percentage of cells traversing the membrane relative to the initial seeding density.

### Cell Invasion Assay

Cell invasion was assessed using a 96-well basement membrane invasion assay with 8 µm pore size (Abcam, ab235697). DMS273 parental and liver-metastatic derivatives 138 and 139 were cultured to approximately 80% confluence and serum-starved for 18-24 hours before the assay. Cells, 5 x 10⁴ per condition, were suspended in serum-free medium and seeded into Boyden chambers pre-coated with basement membrane extract, with chemoattractant-containing medium placed in the lower chamber.

Following incubation for 48 hours at 37°C in 5% CO₂, non-invasive cells were removed from the upper chamber. Invaded cells were collected from the lower chamber, dissociated, stained with fluorescent dye, and quantified by fluorescence measurement at Ex/Em = 530/590 nm. Fluorescence values were converted to cell numbers using a standard curve generated by serial dilution of known cell densities. Invasion was expressed as the percentage of cells traversing the matrix-coated membrane relative to the initial seeding density.

### Cell Line Bulk RNA Sequencing

Liver metastasis-derived cell lines from patient or animal models, approximately 0.5 million cells per sample, were harvested, washed with PBS, and processed for total RNA extraction using the RNAqueous-4PCR DNA-Free RNA Isolation Kit (Invitrogen, Cat. No. AM1914). Three technical replicates were generated per cell line. RNA quality and integrity were verified using the Agilent 4150 TapeStation System with a standard sensitivity RNA assay. Libraries were prepared using poly(A) enrichment and sequenced on NovaSeq PE150 to approximately 6 Gb per sample.

Paired-end reads were aligned to the human reference genome hg38 using STAR (v2.7.11b). BAM files were generated, sorted, and indexed using SAMtools (v1.17). Gene-level read counts were extracted using STAR’s --quantMode GeneCounts function and normalized to RPKM values. Differential expression analysis was performed using DESeq2 in R (v4.2.3). Genes with log2 fold change > 1.5 or < -1.5 and adjusted P value < 0.05 were considered significantly differentially expressed. Pathway analysis was conducted using Ingenuity Pathway Analysis, with differentially enriched pathways filtered based on absolute z-score > 1 and -log10 P value > 1.3, and prioritized by z-score.

### Western Blotting

Cells, approximately 2 million per condition, were washed with cold PBS and lysed in 250 µL of 2x Laemmli sample buffer (Bio-Rad, Cat. No. 1610737) supplemented with 5% 2-mercaptoethanol (Sigma, Cat. No. M3148-100ML). Lysates were sonicated using five cycles of 30 seconds on/off at 4°C, vortexed, boiled for 5 minutes, and centrifuged at 11,000 x g for 5 minutes at 4°C. Supernatants, 10 µL per sample, were resolved by SDS-PAGE using 10% gels for total protein and 15% gels for histones, followed by transfer to PVDF membranes (Millipore, Cat. No. IPVH304F0).

Membranes were blocked in 5% non-fat dry milk (Bio-Rad) in 0.1% TBS-T, composed of 10 mM Tris-HCl pH 7.6, 150 mM NaCl, and 0.1% Tween-20, for 1 hour at room temperature. Membranes were incubated overnight at 4°C with primary antibodies diluted 1:1000 in 1% BSA in 0.1% TBS-T. After washing, membranes were incubated with HRP-conjugated secondary antibodies diluted 1:10,000 in 5% milk for 1 hour at room temperature. Signal was detected using Immobilon Western or SuperSignal West Femto Chemiluminescent substrates.

### Histone Extraction from Cell Lines

Histones were isolated using a low-concentration acid extraction protocol following nuclear isolation. Cells were washed twice with PBS, and nuclei were extracted using the EpiQuik Nuclear Extraction Kit (Epigentek, Cat. No. OP-0002-1) at 4°C. Nuclei pellets were washed twice with lysis buffer to remove debris. Histone extraction was performed by resuspending the nuclei pellet in 0.3 mL of 0.2 M H₂SO₄, approximately six volumes, followed by vigorous vortexing at 4°C for 30 minutes. The suspension was centrifuged at 16,200 rpm for 10 minutes at 4°C, and the supernatant was transferred to a new tube for protein precipitation at -20°C overnight.

Precipitated histones were collected by centrifugation and washed sequentially with 50 mM HCl in acetone and then with chilled acetone alone. The final histone pellet was air-dried and resuspended in 0.1% β-mercaptoethanol in H₂O, then stored at -20°C. Histone purity was confirmed by 15% SDS-PAGE before western blot analysis.

### Cell Fractionation

Cell fractionation was performed with modifications based on a previously published protocol^62^. Briefly, cells were harvested and resuspended in 200 µL Buffer A composed of 10 mM HEPES pH 8.0, 10 mM KCl, 1.5 mM MgCl₂, 0.34 M sucrose, 10% glycerol, 1 mM DTT, 1 mM PMSF, 0.1% Triton X-100, and 1x phosphatase and protease inhibitors. Cells were incubated on ice for 5 minutes. The lysate was centrifuged at 1300 x g for 4 minutes at 4°C to separate the cytoplasmic supernatant from the nuclear pellet. The nuclear pellet was washed twice with Buffer A, resuspended in 200 µL Buffer B composed of 3 mM EDTA, 0.2 mM EGTA, 1 mM DTT, 1 mM PMSF, and 1x phosphatase and protease inhibitors, and incubated on ice for 30 minutes.

The nuclear lysate was centrifuged at 1700 x g for 4 minutes at 4°C to separate the soluble nuclear fraction from the chromatin pellet, which was washed three times in Buffer B. To obtain a cleared cytoplasmic fraction, the cytoplasmic supernatant was further centrifuged at 14,000 x g for 15 minutes at 4°C, and the resulting supernatant was retained. All fractions were subjected to downstream analysis by western blotting.

### Chromatin Immunoprecipitation and qPCR (ChIP-qPCR)

Chromatin immunoprecipitation assays were performed using the SimpleChIP Plus Enzymatic Chromatin IP Kit (Cell Signaling Technology, Cat. No. 9005), according to the manufacturer’s protocol. Briefly, 4 million cells were crosslinked with 1% formaldehyde for 10 minutes, quenched with 125 mM glycine for 5 minutes, and lysed in Buffer A (CST #7006). Nuclei were pelleted, washed in Buffer B (CST #7007), and digested with 0.5 µL micrococcal nuclease (CST #10011) at 37°C for 20 minutes with intermittent mixing to generate approximately 150–900 bp fragments. Digestion was stopped with 0.5 M EDTA, and nuclei were pelleted and lysed in ChIP Buffer (CST #7008).

Chromatin was sonicated using a Bioruptor (Diagenode) and pre-cleared with Dynabeads Protein A (Thermo Fisher). Immunoprecipitation was performed overnight at 4°C with Dynabeads pre-bound to specific antibodies. Beads were sequentially washed with low-salt buffer, high-salt buffer composed of ChIP buffer plus 75 µL 5 M NaCl, and TE buffer. Bound chromatin was eluted twice with 100 µL elution buffer (CST #7009), and crosslinks were reversed by overnight incubation at 65°C with 0.1 mg/mL RNase A, 0.3 M NaCl, and 0.2 mg/mL proteinase K. DNA was purified using QIAquick PCR columns (Qiagen, Cat. No. 28104).

Quantitative PCR was performed on a Bio-Rad CFX Connect Real-Time PCR System using SYBR Green Supermix (Bio-Rad, Cat. No. 1725271). Primers targeting H3K27Ac enrichment at the promoters of HNF1A, HNF4A, Aldolase B, and SOX9, and HIF1α binding at the ACLY hypoxia response element site, were used. Primer sequences are listed in the Key Resources Table.

### Lactate Quantification

Lactate concentrations in cell culture media were measured using the L-Lactic Acid Colorimetric Assay Kit (Elabscience, Cat. No. E-BC-K044-S). DMS273 parental cells, liver-metastatic clones 137, 138, and 139, and HHL5 hepatocytes were cultured in DMEM supplemented with 10% FBS. After overnight incubation, cells were starved for 40 minutes in DMEM containing 10% charcoal-stripped serum without a carbon source. Cells were then incubated for 2 hours in DMEM supplemented with 10% FBS and 4.5 g/L glucose. Media and cell lysates were harvested, cleared of debris by centrifugation, and analyzed according to the manufacturer’s instructions. Lactate levels were normalized to total cellular protein content.

### Subcellular Fractionation and Acetyl-CoA Quantification

SCLC cells were fractionated using the Cell Fractionation Kit (Cell Signaling Technology, Cat. No. 9038), yielding whole-cell lysate, cytosolic, and nuclear fractions. Protein concentrations were quantified using the Qubit Protein Assay Kit (Invitrogen, Cat. No. Q33212). Twenty micrograms of protein per fraction were used to quantify acetyl-CoA levels using an ELISA-based Acetyl-CoA Assay Kit (Elabscience, Cat. No. E-EL-0125), according to the manufacturer’s protocol.

### Nuclear HAT Activity Assay

Histone acetyltransferase activity in nuclear lysates was measured using the EpiQuik HAT Activity/Inhibition Assay Kit (Epigentek, Cat. No. P-4003-96). DMS273 parental cells and metastatic clones 137, 138, and 139 were cultured in DMEM supplemented with 10% FBS. Nuclear proteins were extracted using the EpiQuik Nuclear Extraction Kit (Epigentek, Cat. No. OP-0002-1) and quantified with the Qubit assay. HAT activity was measured using 20 µg of nuclear protein, normalized to total protein content, and reported as relative enzymatic activity.

### Nuclear HDAC Activity Assay

Nuclear histone deacetylase activity was quantified using the HDAC Activity Assay Kit (Epigentek, Cat. No. P-4034-96), according to the manufacturer’s protocol. Nuclear lysates were prepared and quantified as described above. A total of 20 µg nuclear protein was used per reaction, and HDAC activity was normalized to protein concentration.

### Glucose Uptake by Flow Cytometry

Glucose uptake was assessed in SCLC cell lines CRL9609, RA22#12 parental, RA22#6, and #404 liver metastasis using 2-NBDG, a fluorescent glucose analog. Cells, 2 x 10⁵ per condition, were plated in DMEM GlutaMAX overnight and then glucose-starved for 40 minutes in DMEM with 10% charcoal-stripped serum. Cells were incubated with 50 µg/mL 2-NBDG in DMEM for 2 hours at 37°C, harvested, washed with PBS, and analyzed using a BD FACSAria Fusion flow cytometer. Fluorescence data were processed with BD FACSDiva software v9.01.

### Real-Time Metabolic Assays

Oxygen consumption rate and extracellular acidification rate were measured using a Seahorse XF96 Analyzer (Agilent). DMS273 parental and liver-metastatic derivatives 137, 138, and 139 were seeded at 40,000 cells per well. After monolayer formation, cells were incubated overnight in DMEM GlutaMAX. Before the assay, cells were washed and incubated for 1 hour in XF assay medium (Agilent, Cat. No. 103575-100) without glucose, glutamine, pyruvate, or serum.

Basal oxygen consumption rate was measured, followed by sequential injections of oligomycin, FCCP, and rotenone/antimycin A as needed. Oxygen consumption rate (OCR) and extracellular acidification rate (ECAR) were measured under varying nutrient conditions, including glucose alone, glucose plus glutamine plus pyruvate, or glutamine alone, and normalized to seeding density.

### Total metabolite and flux analysis using 13C-Glc

Metabolic flux analysis using 13C-Glc was performed by Human Metabolome Technologies, Inc. Three biological replicates each of DMS273 parental cells and the liver-adapted derivatives 137, 138, and 139 were used for this experiment. Cells were grown in medium containing unlabeled pyruvate and glutamine, and 13C-labeled glucose (13C-Glc). Samples were prepared according to the service provider’s guidelines from 5 × 10^6^ cells per replicate and resuspended in 50 μl ultrapure water before measurement. The samples were analyzed using capillary electrophoresis time-of-flight mass spectrometry (CE-TOFMS, Agilent Technologies) in two modes to detect both anionic and cationic metabolites. Detected peaks were then extracted using MasterHands ver. 2.17.1.11 to obtain m/z, migration time (MT), and peak area. Putative metabolites were assigned based on HMT’s target library and their isotopic ions using m/z and MT. Absolute quantitation was performed for the total amount of each detected metabolite.

### Correlative FIB-SEM

For sample preparation, DMS273-RFP cells and HHL5-GFP cells were cultured on 35 mm grid-patterned glass-bottom plates for 24 hours (MatTek). Cells were imaged by fluorescence microscopy and then immediately fixed with Karnovsky’s fixative, composed of 2% paraformaldehyde and 2% glutaraldehyde in 0.1 M sodium cacodylate, for approximately 2 hours at room temperature. Grid coordinates for specific regions of interest were imaged by transmitted light.

Co-cultured cells were post-fixed in 2% osmium tetroxide and 1.5% potassium ferricyanide in 0.1 M sodium cacodylate for 1 hour at room temperature, washed five times with ultrapure water, and stained with 1% aqueous uranyl acetate overnight at 4°C. After washing with water, samples were treated with lead aspartate at 65°C for 30 minutes and washed again. Cells were dehydrated through graded ethanol, including 35%, 50%, 70%, 95%, and five changes of 100% ethanol, for 10 minutes each. Samples were infiltrated with increasing concentrations of Polybed 812 resin with ethanol at resin:ethanol ratios of 1:3, 1:2, 3:1, and 100% resin, embedded in 100% degassed resin, and polymerized at 65°C for 48 hours. Glass coverslips were removed from the resin by heat shock. Resin specimens containing regions of interest identified by light microscopy were cut using a jeweler’s saw and mounted on pin stubs, silver painted, and carbon-coated to approximately 6 nm using a Leica coater.

Cell regions of interest previously identified by light microscopy were located by SEM imaging on a Zeiss Crossbeam 550 using the gridded pattern implanted in the resin. The FIB sample preparation workflow was initiated using the Fibics ATLAS3D sample preparation workflow. A 1 µm platinum pad was laid over the area containing target cells. Tracking and autofocus lines were milled into the platinum surface and coated with a 1 µm carbon pad. After pad completion, a coarse trench was milled using a 30 nA FIB beam, stopping just short of the platinum pad’s leading edge. The trench face was then fine-polished using a 3 nA beam.

SEM imaging parameters were set at 1.5 kV accelerating voltage with a 1.1 nA beam current, pixel resolution of 5 nm, slice thickness of 10 nm, and EsB detector grid voltage of 790 V. Raw image stacks were processed through registration, inversion, and binning to produce volumetric reconstructions at 10 nm isotropic voxels, following established protocols^63^. Mitochondria and nuclei were segmented using deep-learning panoptic segmentation models implemented in the empanada plugin for napari^44^ from image volumes binned by two. Three-dimensional visualization of organelles was performed in Dragonfly 3D, version 2025.1.

### Global Proteomics Analysis

Cell pellets were resuspended in 200 µL EasyPep buffer (Thermo, Cat. No. A45735) with 1 µL nuclease and PhosSTOP. Protein concentration was estimated using BCA assay, and 100 µg of each sample was combined with 50 µL reduction solution, TCEP at 16 mM, and 50 µL alkylation solution, chloroacetamide at 32 mM. Five micrograms of Trypsin/LysC (Thermo, Cat. No. 90051) was added to each sample, followed by overnight digestion at 37°C with shaking at 500 rpm.

The next day, 200 µg TMTpro label (Thermo, Cat. No. A52045) was added to each sample and incubated for 1 hour at 25°C. Reactions were quenched with 50 µL of 5% hydroxylamine and 20% formic acid for 10 minutes. Samples were combined and cleaned using the EasyPep Maxi Kit (Thermo, Cat. No. A45734), according to the manufacturer’s instructions. Following elution, samples were combined and dried in a speed-vac.

The sample was resuspended in 50 µL of 0.1% formic acid, and 5 µL was loaded onto a Dionex U3000 RSLC coupled to an Orbitrap Eclipse mass spectrometer equipped with FAIMS and an EasySpray ion source. Solvent A was 0.1% formic acid, and solvent B was 0.1% formic acid in 80% acetonitrile. The gradient pump was run at 300 µL/min with an LC gradient of 5-7% B for 1 minute, 7-30% B for 140 minutes, 30-50% B for 35 minutes, 50-95% B for 4 minutes, 95% B for 7 minutes, followed by re-equilibration of the column at 5% B for 17 minutes.

The mass spectrometer was run in TopSpeed mode with four FAIMS compensation voltages, either 45, 55, 65, and 75 or 50, 60, 70, and 80, with 1-second cycle time for each compensation voltage. For each compensation voltage, the spray voltage was set at 2200 V and the ion transfer tube temperature at 300°C. MS1 scans were acquired in the Orbitrap at 120,000 resolution, AGC of 4e5, and maximum injection time of 50 ms over a mass range of 375-1600 m/z. MS2 scans were acquired in the Orbitrap using the TurboTMT method with a resolution of 15,000, intensity threshold of 2.5e4, normalized AGC target of 250%, HCD energy of 38%, isolation window of 0.4 m/z, and charges of 2-5 selected for MS2. Monoisotopic precursor selection, Easy-IC for internal calibration, and advanced peak determination were enabled.

MS files were searched in Proteome Discoverer 2.4 using the SequestHT node. Data were searched against the UniProt Human database, February 2020 reviewed canonical set, using full trypsin digestion with a maximum of two missed cleavages, minimum peptide length of six, MS1 mass tolerance of 10 ppm, and MS2 tolerance of 0.02 Da. Variable modification included oxidation of methionine (+15.995), and static modifications included carbamidomethylation of cysteine (+57.021) and TMTpro labeling of lysine and peptide N-termini (+304.207). Percolator was used for false-discovery-rate analysis, and TMTpro reporter ions were quantified using the Reporter Ion Quantifier node and normalized on total peptide intensity of each channel.

### Animal Studies

All animal studies were conducted in accordance with protocols approved by the Animal Care and Use Committee. All procedures were performed in accordance with the guidelines outlined in the *Guide for the Care and Use of Laboratory Animals*.

### Intracardiac Injection Model

To generate systemic metastases, including liver and brain metastases, four- to six-week-old male and female NSG mice, NOD.Cg-Prkdc^scid Il2rg^tm1Wjl/SzJ, The Jackson Laboratory #005557, were anesthetized and injected with 5 x 10⁵ DMS273 cells into the left cardiac ventricle under ultrasound guidance. Tumor burden was monitored weekly using magnetic resonance imaging and bioluminescence imaging.

### Orthotopic Lung Tumor Model

Orthotopic small cell lung tumors were established by injecting 1 x 10⁵ RA22#12 patient-derived SCLC cells into the left lung lobe of anesthetized NSG mice. Cells were suspended in PBS or a 1:1 PBS–Matrigel mixture. A left thoracotomy was performed to access the injection site. The incision was closed with sutures or tissue adhesive. Post-operative analgesia was provided according to Animal Care and Use Committee guidelines. Tumor burden was assessed weekly using bioluminescence imaging for luciferase-expressing cells, and lungs and livers were harvested at endpoint or upon signs of morbidity.

### Intrasplenic Injection Model

NSG mice were anesthetized with isoflurane, and the thoracic and peritoneal regions were shaved and disinfected with three alternating rounds of betadine and ethanol. Mice were placed on a heating pad to maintain body temperature and given pre-operative subcutaneous fluids, 1 cc Normosol-R or normal saline pre-warmed to 37°C. Through a left lateral flank incision of approximately 1 cm, the spleen was exteriorized, and a tumor cell suspension containing 5 x 10⁵ RA22#4 tumor cells in 50 µL PBS was injected using a 27-gauge needle. Visible paling of the spleen and lack of bleeding were used as criteria for successful inoculation.

The cell suspension was allowed to enter the portal circulation over 5 minutes, after which the spleen was removed to prevent growth of tumor cells in the spleen. The spleen was raised to expose vessels attaching it to the pancreas. The vessels between the spleen and pancreas were cut by cauterization. The spleen was removed, and the peritoneal cavity was sutured using polyglycolic acid, Dexon 5-0. The skin was closed with either sutures or wound clips. A few drops of bupivacaine were placed in the incision site before closure to alleviate postoperative pain. Mice were allowed to recover from anesthesia on a warming tray and then placed in clean cages. One dose of sustained-release buprenorphine was used as an analgesic. Following surgery, mice were monitored twice the same day and then daily until complete recovery. Wound clips or sutures were removed approximately 7–10 days after surgery.

### In Vivo Imaging

For bioluminescence imaging, mice were imaged using an IVIS Spectrum system (Revvity, Waltham, MA) in both dorsal and ventral positions. Imaging parameters included excitation filter blocked, emission filter open, f/stop 1, medium binning of 8 x 8, and automatic exposure from 1 to 120 seconds. Living Image software v4.7.4 was used for image acquisition and analysis. Tumor burden was quantified using fixed-size regions of interest, rectangular for liver in ventral images and circular for brain in dorsal images.

For magnetic resonance imaging, imaging was initiated two weeks post-injection using a 3T clinical scanner, Intera-Achieva 3T, Philips, Netherlands, with a custom-built volume receive array coil (Lambda Z Technologies, MD) for simultaneous imaging of three mice. T2-weighted turbo spin echo sequences were acquired in the coronal plane from brain to abdomen using the following parameters: field of view 78 x 160 mm, repetition time 5880 ms, echo time 45 ms, resolution 0.18 x 0.18 mm², and slice thickness 0.5 mm. Spectral presaturation with inversion recovery fat suppression was used to enhance soft tissue contrast.

### Antibodies, Reagents, Primers, Plasmids, and shRNA Sequences

All antibodies, primer sequences, plasmid vectors, and shRNA constructs used in this study are detailed in the Key Resources Table.

### Statistical Analysis

Comparisons between two groups were performed using two-sided Wilcoxon rank-sum tests or unpaired two-tailed Student’s t-tests, as indicated. For multi-group comparisons, one-way ANOVA with Tukey post hoc testing was used where appropriate. Quantitative pathology data from HALO were exported into GraphPad Prism 10.4.1 for statistical analysis. Survival analyses were performed using Cox proportional hazards models and Kaplan-Meier estimates. P values < 0.05 were considered statistically significant.

## References

1. Fidler IJ, Kripke ML. The challenge of targeting metastasis. Cancer Metastasis Rev. 2015;34(4):635–641.

2. Gerstberger S, Jiang Q, Ganesh K. Metastasis. Cell. 2023;186(8):1564–1579.

3. Massague J, Obenauf AC. Metastatic colonization by circulating tumour cells. Nature. 2016;529(7586):298–306.

4. Mielgo A, Schmid MC. Liver Tropism in Cancer: The Hepatic Metastatic Niche. Cold Spring Harb Perspect Med. 2020;10(3).

5. Tsilimigras DI, Brodt P, Clavien PA, et al. Liver metastases. Nat Rev Dis Primers. 2021;7(1):27.

6. Fazilaty H, Basler K. Reactivation of embryonic genetic programs in tissue regeneration and disease. Nature Genetics. 2023;55(11):1792–1806.

7. Ganesh K, Basnet H, Kaygusuz Y, et al. L1CAM defines the regenerative origin of metastasis-initiating cells in colorectal cancer. Nat Cancer. 2020;1(1):28–45.

8. Moorman A, Benitez EK, Cambulli F, et al. Progressive plasticity during colorectal cancer metastasis. Nature. 2025;637(8047):947–954.

9. Househam J, Heide T, Cresswell GD, et al. Phenotypic plasticity and genetic control in colorectal cancer evolution. Nature. 2022;611(7937):744–753.

10. Laughney AM, Hu J, Campbell NR, et al. Regenerative lineages and immune-mediated pruning in lung cancer metastasis. Nat Med. 2020;26(2):259–269.

11. Bala P, Rennhack JP, Aitymbayev D, et al. Aberrant cell state plasticity mediated by developmental reprogramming precedes colorectal cancer initiation. Sci Adv. 2023;9(13):eadf0927.

12. McDonald OG, Li X, Saunders T, et al. Epigenomic reprogramming during pancreatic cancer progression links anabolic glucose metabolism to distant metastasis. Nature Genetics. 2017;49(3):367–376.

13. Pomerantz MM, Qiu X, Zhu Y, et al. Prostate cancer reactivates developmental epigenomic programs during metastatic progression. Nat Genet. 2020;52(8):790–799.

14. Abbott KL, Subudhi S, Ferreira R, et al. Nutrient requirements of organ-specific metastasis in breast cancer. Nature. 2026;649(8099):1292–1301.

15. Thomas A, Mohindroo C, Giaccone G. Advancing therapeutics in small-cell lung cancer. Nat Cancer. 2025.

16. Rudin CM, Brambilla E, Faivre-Finn C, Sage J. Small-cell lung cancer. Nat Rev Dis Primers. 2021;7(1):3.

17. Thomas A, Pattanayak P, Szabo E, Pinsky P. Characteristics and Outcomes of Small Cell Lung Cancer Detected by CT Screening. Chest. 2018;154(6):1284–1290.

18. Nakazawa K, Kurishima K, Tamura T, et al. Specific organ metastases and survival in small cell lung cancer. Oncol Lett. 2012;4(4):617–620.

19. Fan L, Lin Y, Fu Y, Wang J. Small cell lung cancer with liver metastases: from underlying mechanisms to treatment strategies. Cancer Metastasis Rev. 2024;44(1):5.

20. Cai H, Wang H, Li Z, Lin J, Yu J. The prognostic analysis of different metastatic patterns in extensive-stage small-cell lung cancer patients: a large population-based study. Future Oncol. 2018;14(14):1397–1407.

21. Kagohashi K, Satoh H, Ishikawa H, Ohtsuka M, Sekizawa K. Liver metastasis at the time of initial diagnosis of lung cancer. Med Oncol. 2003;20(1):25–28.

22. Ko J, Winslow MM, Sage J. Mechanisms of small cell lung cancer metastasis. EMBO Mol Med. 2021;13(1):e13122.

23. Disibio G, French SW. Metastatic patterns of cancers: results from a large autopsy study. Arch Pathol Lab Med. 2008;132(6):931–939.

24. Chen HZ, Bonneville R, Paruchuri A, et al. Genomic and Transcriptomic Characterization of Relapsed SCLC Through Rapid Research Autopsy. JTO clinical and research reports. 2021;2(4):100164.

25. Chan JM, Quintanal-Villalonga A, Gao VR, et al. Signatures of plasticity, metastasis, and immunosuppression in an atlas of human small cell lung cancer. Cancer Cell. 2021;39(11):1479–1496 e1418.

26. Kawasaki K, Salehi S, Zhan YA, et al. FOXA2 promotes metastatic competence in small cell lung cancer. Nat Commun. 2025;16(1):4865.

27. Liu Q, Zhang J, Guo CC, et al. Proteogenomic characterization of small cell lung cancer identifies biological insights and subtype-specific therapeutic strategies. Cell. 2024;187(1).

28. George J, Lim JS, Jang SJ, et al. Comprehensive genomic profiles of small cell lung cancer. Nature. 2015;524(7563):47–53.

29. Chen WS, Alshalalfa M, Zhao SG, et al. Novel RB1-Loss Transcriptomic Signature Is Associated with Poor Clinical Outcomes across Cancer Types. Clin Cancer Res. 2019;25(14):4290–4299.

30. Segal JM, Kent D, Wesche DJ, et al. Single cell analysis of human foetal liver captures the transcriptional profile of hepatobiliary hybrid progenitors. Nature Communications. 2019;10.

31. Tarlow BD, Pelz C, Naugler WE, et al. Bipotential adult liver progenitors are derived from chronically injured mature hepatocytes. Cell Stem Cell. 2014;15(5):605–618.

32. Li W, Yang L, He Q, et al. A Homeostatic Arid1a-Dependent Permissive Chromatin State Licenses Hepatocyte Responsiveness to Liver-Injury-Associated YAP Signaling. Cell Stem Cell. 2019;25(1):54–68 e55.

33. Yanger K, Zong Y, Maggs LR, et al. Robust cellular reprogramming occurs spontaneously during liver regeneration. Genes Dev. 2013;27(7):719–724.

34. Li L, Cui L, Lin P, et al. Kupffer-cell-derived IL-6 is repurposed for hepatocyte dedifferentiation via activating progenitor genes from injury-specific enhancers. Cell Stem Cell. 2023;30(3):283–+.

35. Fares J, Fares MY, Khachfe HH, Salhab HA, Fares Y. Molecular principles of metastasis: a hallmark of cancer revisited. Signal Transduct Target Ther. 2020;5(1):28.

36. Kietzmann T. Metabolic zonation of the liver: The oxygen gradient revisited. Redox Biol. 2017;11:622–630.

37. Deng Z, Fan T, Xiao C, et al. TGF-beta signaling in health, disease, and therapeutics. Signal Transduct Target Ther. 2024;9(1):61.

38. Antoniou A, Raynaud P, Cordi S, et al. Intrahepatic bile ducts develop according to a new mode of tubulogenesis regulated by the transcription factor SOX9. Gastroenterology. 2009;136(7):2325–2333.

39. Kardassis D, Pardali K, Zannis VI. SMAD proteins transactivate the human ApoCIII promoter by interacting physically and functionally with hepatocyte nuclear factor 4. J Biol Chem. 2000;275(52):41405–41414.

40. Chou WC, Prokova V, Shiraishi K, et al. Mechanism of a transcriptional cross talk between transforming growth factor-beta-regulated Smad3 and Smad4 proteins and orphan nuclear receptor hepatocyte nuclear factor-4. Mol Biol Cell. 2003;14(3):1279–1294.

41. Sanchez-Elsner T, Botella LM, Velasco B, Corbi A, Aƫsano L, Bernabeu C. Synergistic cooperation between hypoxia and transforming growth factor-beta pathways on human vascular endothelial growth factor gene expression. J Biol Chem. 2001;276(42):38527–38535.

42. Thakur A, Wong JCH, Wang EY, et al. Hepatocyte Nuclear Factor 4-Alpha Is Essential for the Active Epigenetic State at Enhancers in Mouse Liver. Hepatology. 2019;70(4):1360–1376.

43. Raisner R, Kharbanda S, Jin L, et al. Enhancer Activity Requires CBP/P300 Bromodomain-Dependent Histone H3K27 Acetylation. Cell Rep. 2018;24(7):1722–1729.

44. Conrad R, Narayan K. Instance segmentation of mitochondria in electron microscopy images with a generalist deep learning model trained on a diverse dataset. Cell Syst. 2023;14(1):58–+.

45. Robey IF, Lien AD, Welsh SJ, Baggett BK, Gillies RJ. Hypoxia-inducible factor-1α and the glycolytic phenotype in tumors. Neoplasia. 2005;7(4):324–330.

46. Kim JW, Tchernyshyov I, Semenza GL, Dang CV. HIF-1-mediated expression of pyruvate dehydrogenase kinase: a metabolic switch required for cellular adaptation to hypoxia. Cell Metab. 2006;3(3):177–185.

47. Papandreou I, Cairns RA, Fontana L, Lim AL, Denko NC. HIF-1 mediates adaptation to hypoxia by actively downregulating mitochondrial oxygen consumption. Cell Metab. 2006;3(3):187–197.

48. Denny SK, Yang D, Chuang CH, et al. Nfib Promotes Metastasis through a Widespread Increase in Chromatin Accessibility. Cell. 2016;166(2):328–342.

49. Nabet BY, Hamidi H, Lee MC, et al. Immune heterogeneity in small-cell lung cancer and vulnerability to immune checkpoint blockade. Cancer Cell. 2024.

50. Savchuk S, Gentry K, Wang W, et al. Neuronal-Activity Dependent Mechanisms of Small Cell Lung Cancer Progression. bioRxiv. 2023.

51. Chalabi Hajkarim M, May M, Amin AD, et al. Cellular states associated with metastatic organotropism and survival in patients with pancreatic ductal adenocarcinoma. Nat Genet. 2025.

52. Wolf FA, Angerer P, Theis FJ. SCANPY: large-scale single-cell gene expression data analysis. Genome Biol. 2018;19(1):15.

53. Korsunsky I, Millard N, Fan J, et al. Fast, sensitive and accurate integration of single-cell data with Harmony. Nat Methods. 2019;16(12):1289–1296.

54. Segal JM, Kent D, Wesche DJ, et al. Single cell analysis of human foetal liver captures the transcriptional profile of hepatobiliary hybrid progenitors. Nat Commun. 2019;10(1):3350.

55. Aibar S, Gonzalez-Blas CB, Moerman T, et al. SCENIC: single-cell regulatory network inference and clustering. Nat Methods. 2017;14(11):1083–1086.

56. Badia IMP, Velez Santiago J, Braunger J, et al. decoupleR: ensemble of computational methods to infer biological activities from omics data. Bioinform Adv. 2022;2(1):vbac016.

57. Stuart T, Butler A, Hoffman P, et al. Comprehensive Integration of Single-Cell Data. Cell. 2019;177(7):1888–1902 e1821.

58. Garcia-Alonso L, Holland CH, Ibrahim MM, Turei D, Saez-Rodriguez J. Benchmark and integration of resources for the estimation of human transcription factor activities. Genome Research. 2019;29(8):1363–1375.

59. Browaeys R, Saelens W, Saeys Y. NicheNet: modeling intercellular communication by linking ligands to target genes. Nature Methods. 2020;17(2):159–+.

60. Clayton RF, Rinaldi A, Kandyba EE, et al. Liver cell lines for the study of hepatocyte functions and immunological response. Liver Int. 2005;25(2):389–402.

61. Krishnamurthy M, Dhall A, Sahoo S, et al. Metastatic organotropism in small cell lung cancer. bioRxiv. 2025.

62. Zou L, Cortez D, Elledge SJ. Regulation of ATR substrate selection by Rad17-dependent loading of Rad9 complexes onto chromatin. Genes Dev. 2002;16(2):198–208.

63. Baena V, Conrad R, Friday P, et al. FIB-SEM as a Volume Electron Microscopy Approach to Study Cellular Architectures in SARS-CoV-2 and Other Viral Infections: A Practical Primer for a Virologist. Viruses-Basel. 2021;13(4).

