## Supplementary figures and images for "Hepatobiliary Progenitor-like Reprogramming in Liver Metastases"

### Supplementary Figures S1 through S6

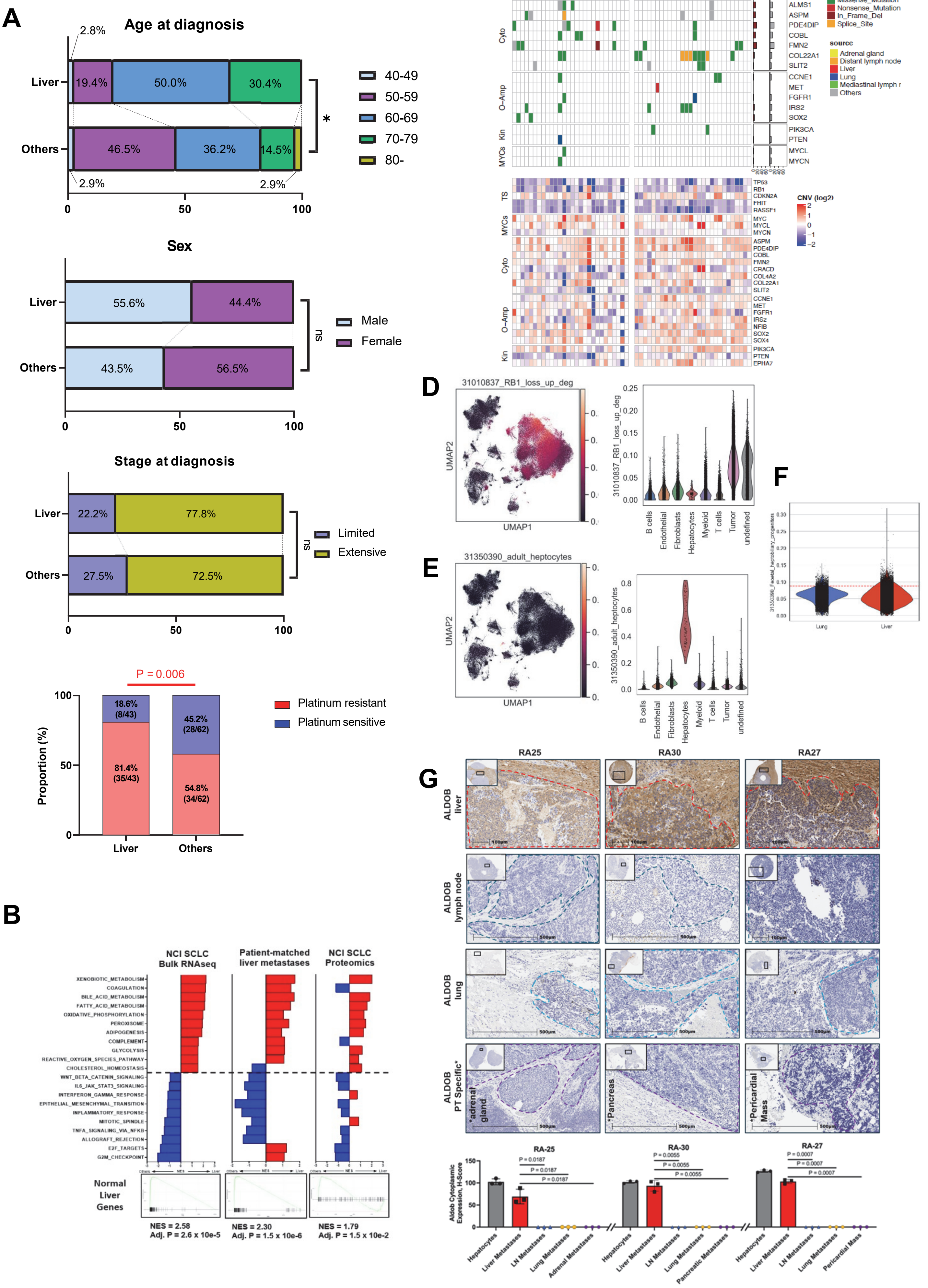

**Fig. S1**

**H**

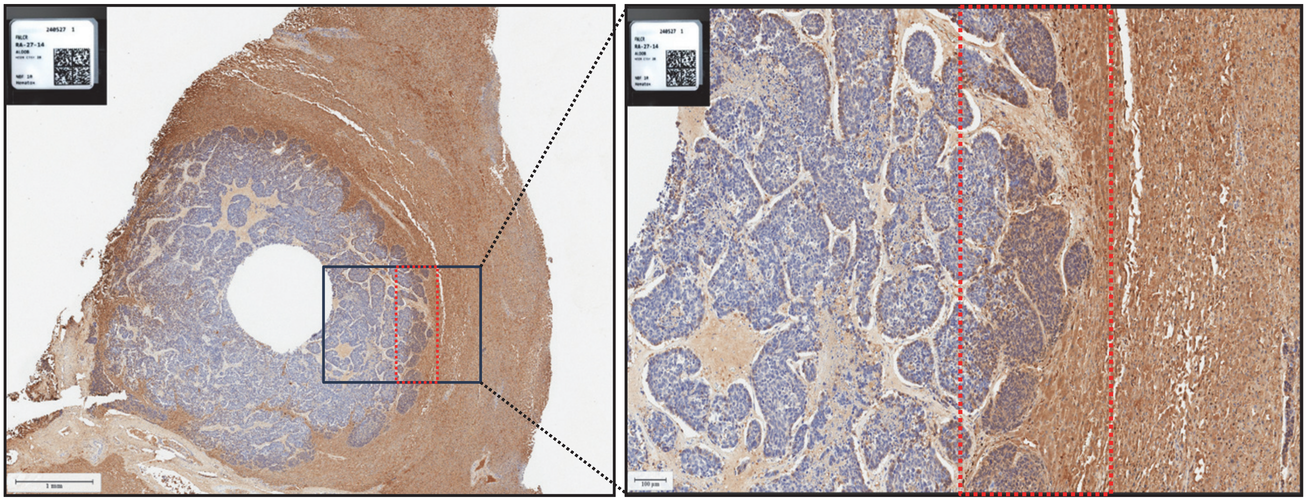

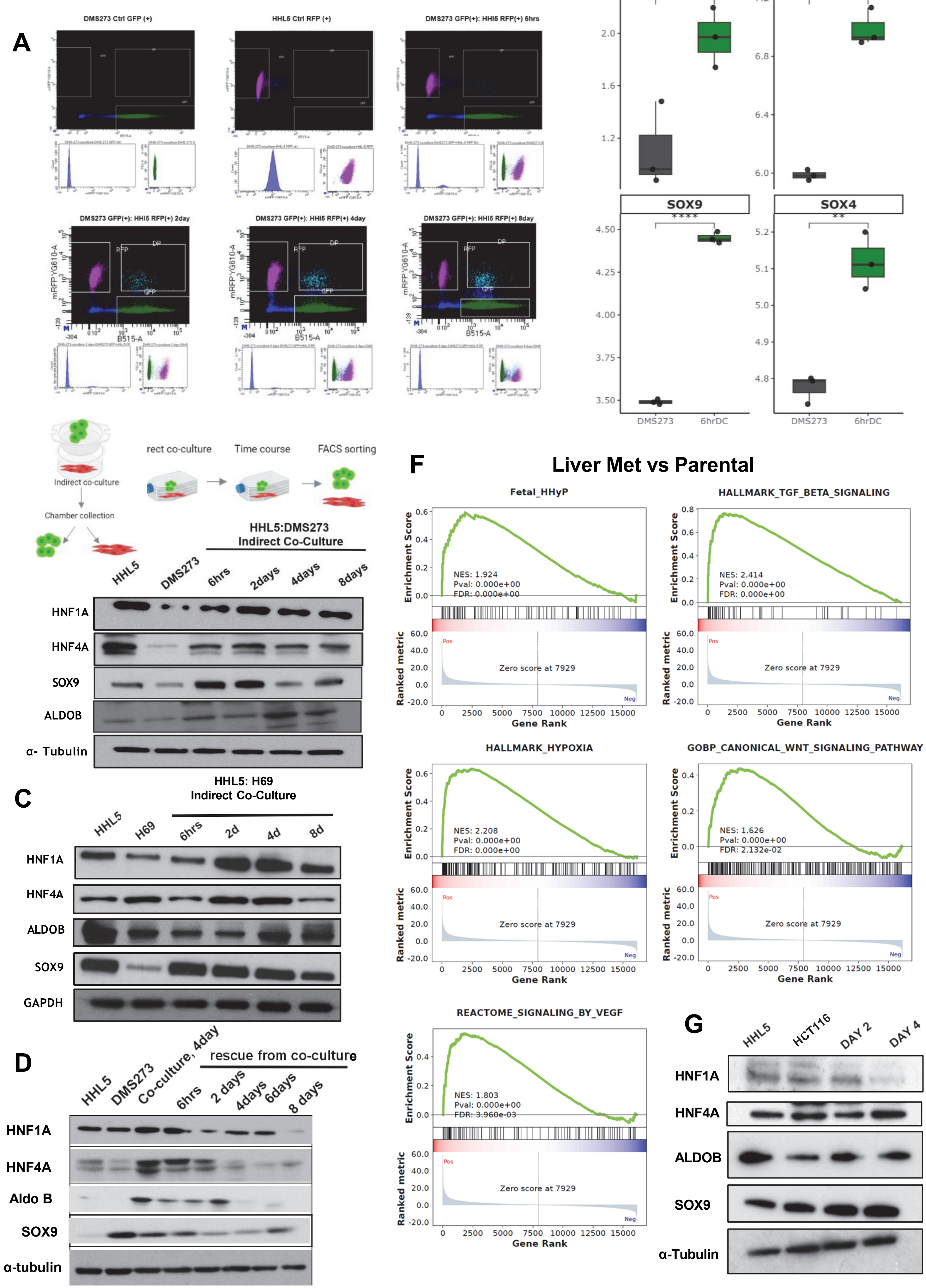

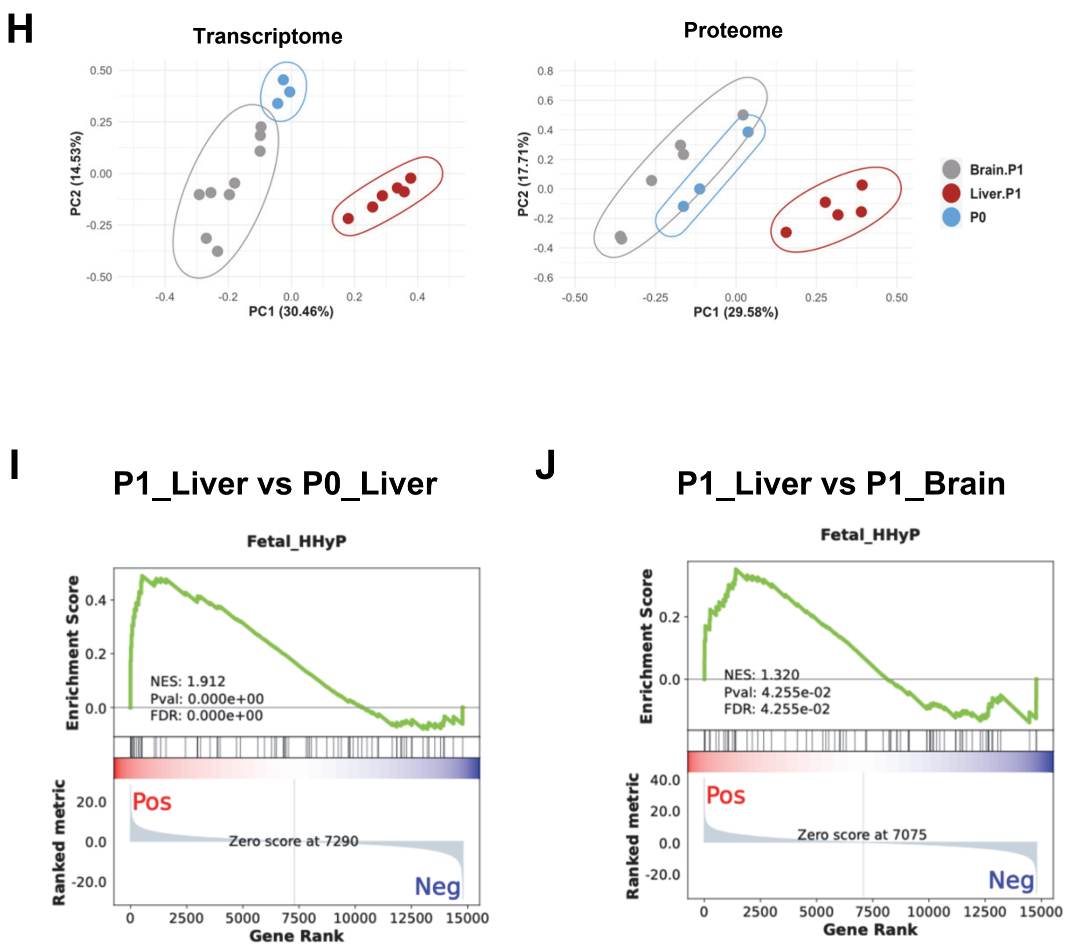

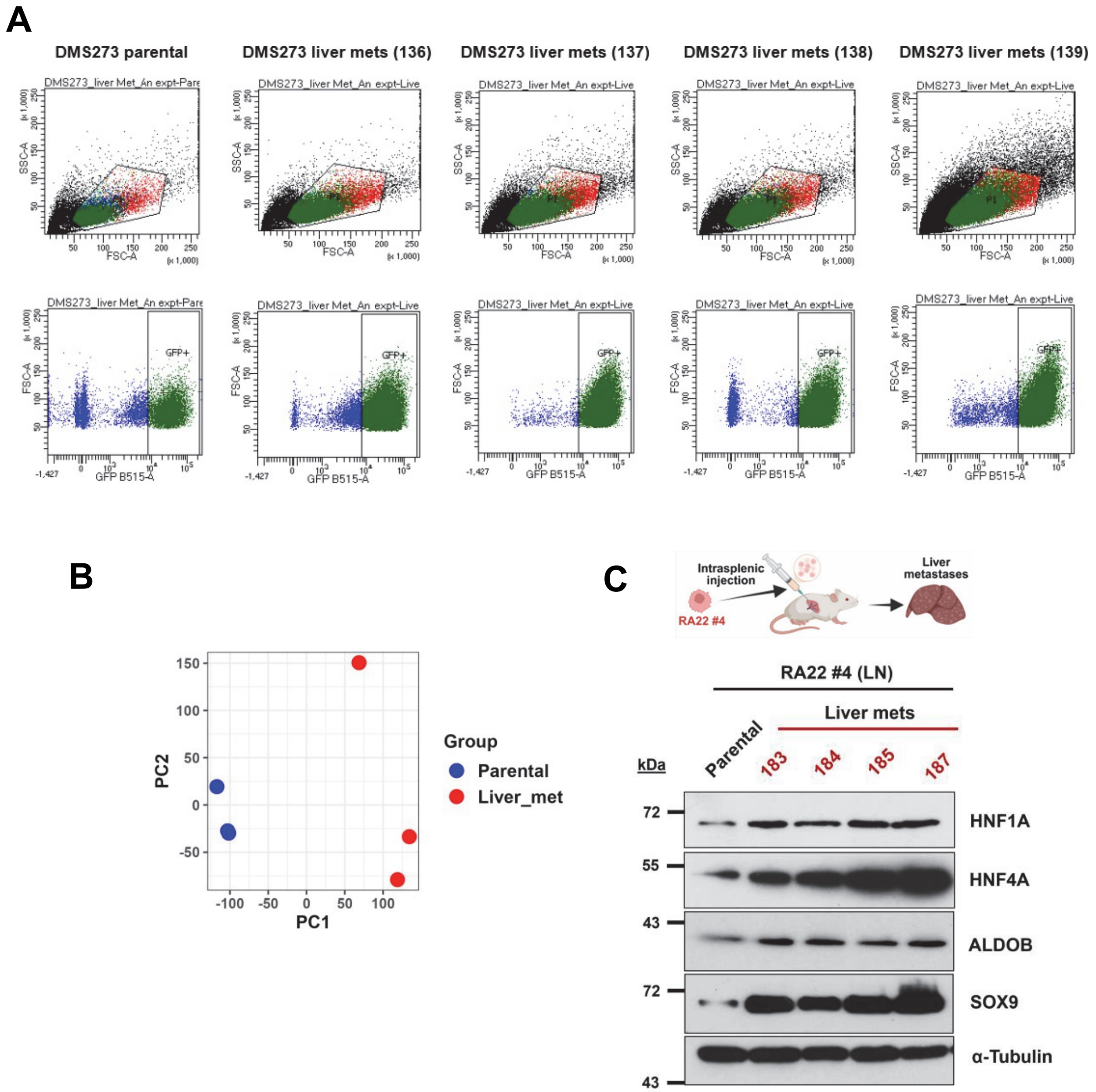

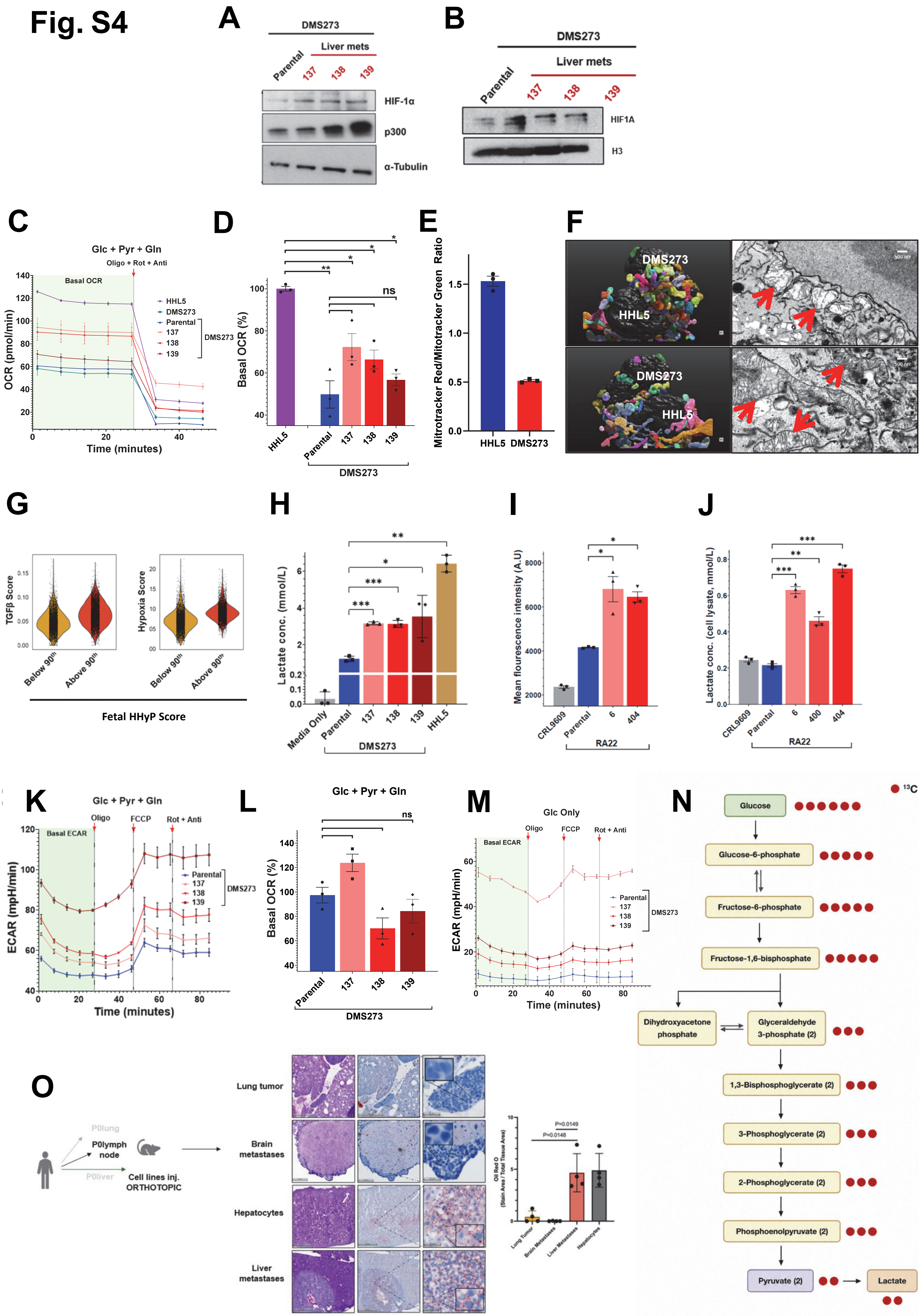

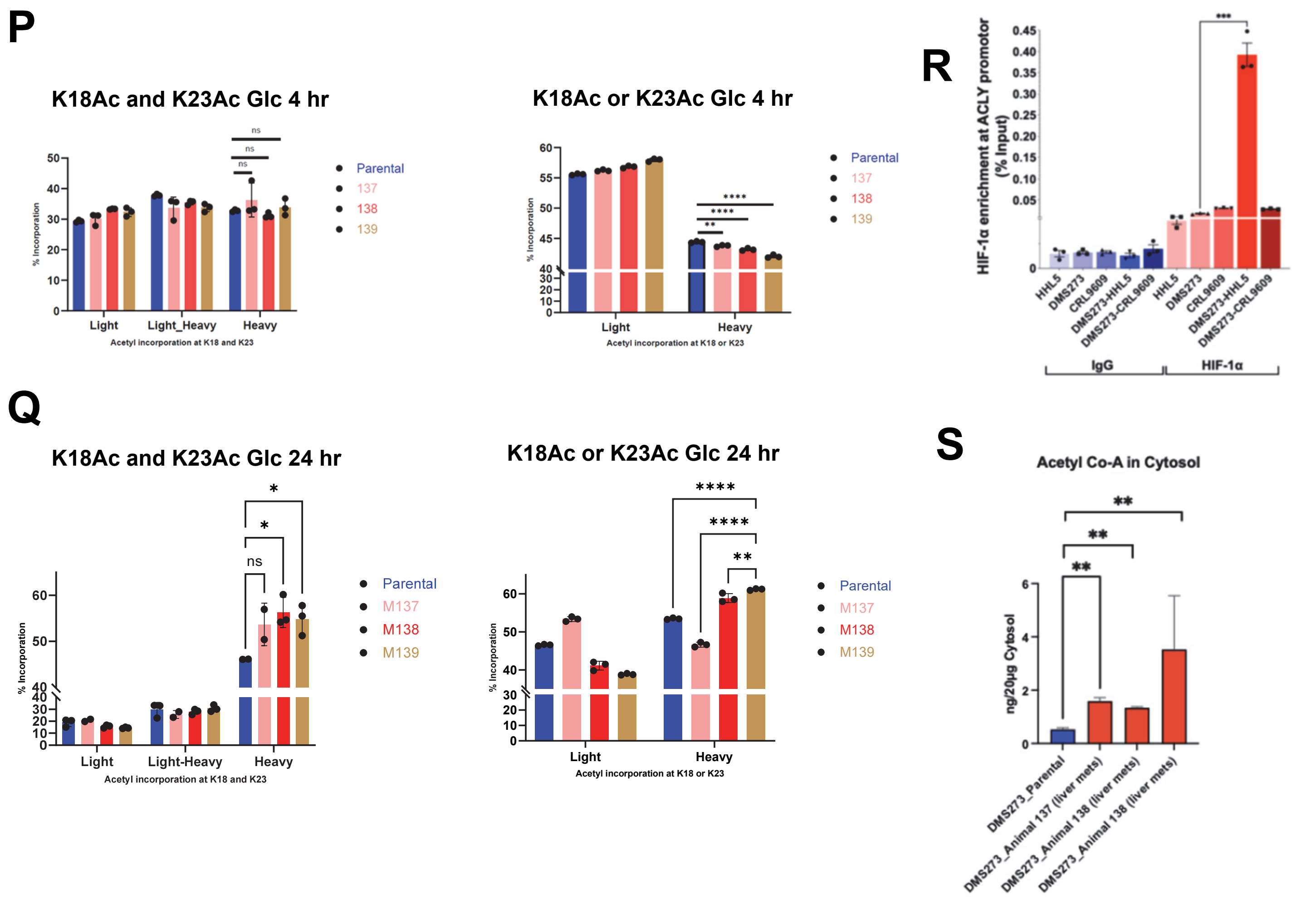

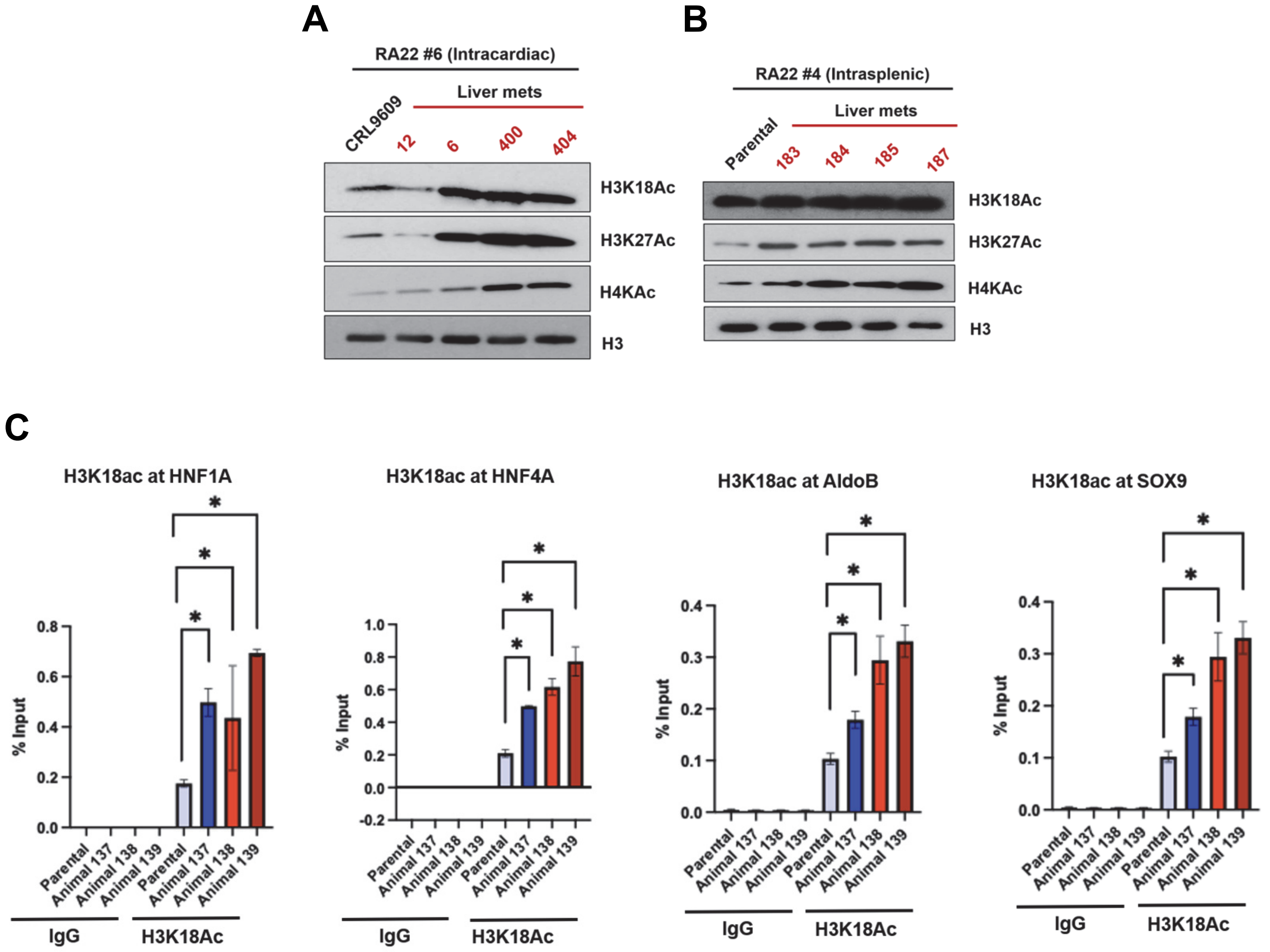

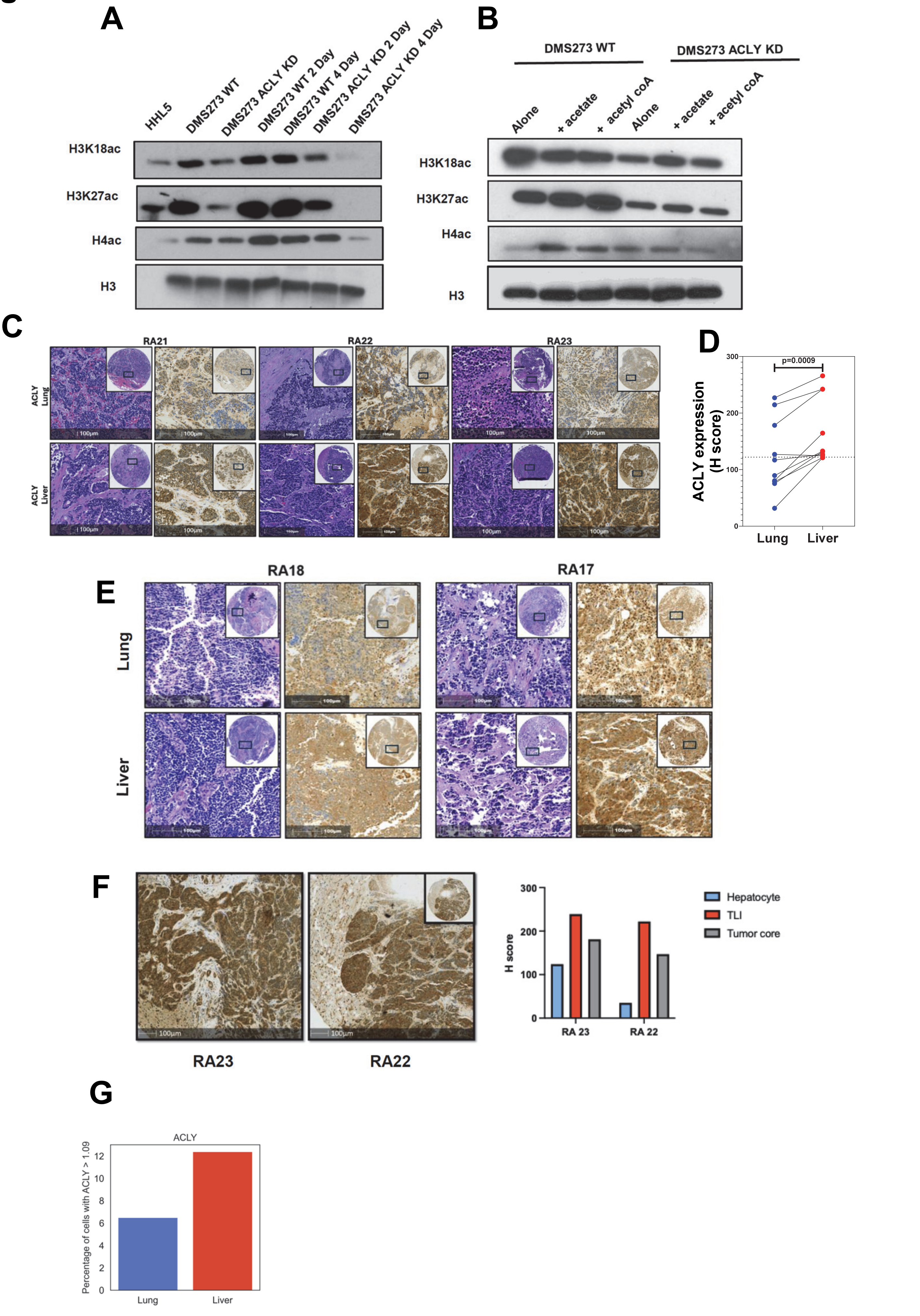
